# pyRBDome: A comprehensive computational platform for enhancing and interpreting RNA-binding proteome data

**DOI:** 10.1101/2023.12.08.570608

**Authors:** Liang-Cui Chu, Niki Christopoulou, Hugh McCaughan, Sophie Winterbourne, Davide Cazzola, Shichao Wang, Ulad Litvin, Salomé Brunon, Patrick J.B. Harker, Iain McNae, Sander Granneman

**Author notes:** These authors contributed equally (Liang-Cui Chu, Niki Christopoulou, Hugh McCaughan). To whom correspondence should be addressed: Sander Granneman.

## Abstract

High-throughput proteomics approaches have revolutionised the identification of RNA-binding proteins (RBPome) and RNA-binding sequences (RBDome) across organisms. Yet the extent of noise, including false-positives, associated with these methodologies, is difficult to quantify as experimental approaches for validating the results are generally low throughput. To address this, we introduce pyRBDome, a pipeline for enhancing RNA-binding proteome data *in silico*. It aligns the experimental results with RNA-binding site (RBS) predictions from distinct machine learning tools and integrates high-resolution structural data when available. Its statistical evaluation of RBDome data enables quick identification of likely genuine RNA-binders in experimental datasets. Furthermore, by leveraging the pyRBDome results, we have enhanced the sensitivity and specificity of RBS detection through training new ensemble machine learning models. pyRBDome analysis of a human RBDome dataset, compared with known structural data, revealed that while UV cross-linked amino acids were more likely to contain predicted RBSs, they infrequently bind RNA in high-resolution structures. This discrepancy underscores the limitations of structural data as benchmarks, positioning pyRBDome as a valuable alternative for increasing confidence in RBDome datasets.

## Introduction

RNA-binding proteins (RBPs) play diverse and crucial roles in gene expression by influencing the structure, function and stability of RNA, both co- and post-transcriptionally. (Holmqvist & Vogel, 2018; Glisovic *et al*, 2008). RBPs have been associated with many human diseases, including neurological disorders, muscular atrophies and cancer (Castello *et al*, 2013). In bacteria, RBPs make key contributions to rapid adaptation to challenging environments, and in pathogens, they control virulence and the capacity for host infections (Christopoulou & Granneman, 2022; Holmqvist & Vogel, 2018). Due to their key functions, considerable efforts are being made to identify RBPs in diverse organisms and to characterise these proteins functionally and structurally. This has inspired the development of several high-throughput methods that capture all proteins interacting with RNA (RBPome). These methods usually involve UV or chemical treatment of cells to create covalent bonds between proteins and direct RNA substrates. This is followed by enrichment of the cross-linked RNA-protein complexes and identification of proteins by quantitative mass spectrometry (MS) (reviewed in (Esteban-Serna *et al*, 2023)). Common approaches for enriching RNA-protein complexes include using oligo(dT) beads to capture proteins cross-linked to polyadenylated RNAs (Castello *et al*, 2012, 2016; Baltz *et al*, 2012; Stenum *et al*, 2023), silica beads that capture all RNAs and cross-linked proteins (Asencio *et al*, 2018; Chu *et al*, 2022; Shchepachev *et al*, 2019; Trendel *et al*, 2019; Beckmann *et al*, 2015; Bae *et al*, 2020) or organic–aqueous phase separation methods that rely on the fact that cross-linked RNAs alter the physiochemical properties of proteins (Queiroz *et al*, 2019; Smith *et al*, 2020; Trendel *et al*, 2019; Urdaneta *et al*, 2019). To identify the cross-linked proteins, purified complexes are treated with ribonucleases and analysed by MS.

These ground-breaking studies have uncovered a plethora of novel RBPs in diverse organisms, many of which contain domains that have never been associated with RNA-binding before. While having a comprehensive list of all RBPs in your favourite organism is tremendously valuable, the next most informative piece of information would be the location of the RNA-binding domains (RBDs) within these proteins (RBDome), as this would allow mechanistic insights into RNA recognition and the design of mutations to dissect the physiological significance of RNA-binding. Although protocols for the global identification of putative RBPs have been optimised for diverse organisms, identifying the amino acid sequences UV cross-linked to RNA (and therefore likely directly bind RNA *in vivo*) in RBPome data is both experimentally and computationally challenging. To identify amino acid-RNA adducts, the cross-linked RNA is chemically or enzymatically digested to make detection of the cross-linking site by MS feasible. However, this digestion is often incomplete, and the heterogeneity in the length and sequence of nucleotide adducts generates variable mass shifts. This dramatically increases the MS/MS search space, making detection of the cross-linking sites using conventional MS data analysis programs unfeasible. To overcome this problem, several experimental computational MS workflows have been developed that either directly detect peptide-RNA conjugates (Kong *et al*, 2017; Kramer *et al*, 2014; Schmidt *et al*, 2012; Trendel *et al*, 2019; Yu *et al*, 2020; Götze *et al*, 2021; Knörlein *et al*, 2022) or identify putative RNA-binding sites (RBSs) by relying on the fact that sequences neighbouring the cross-linked peptides *can* be identified by conventional MS (RBDmap; (Castello *et al*, 2016)), allowing extrapolation of sequences most likely cross-linked to RNA. Recent RBDome methods (RBS-ID and pRBS-ID) utilise hydrofluoride to chemically digest RNAs cross-linked to peptides to a single nucleotide (Bae *et al*, 2020, 2021). This greatly reduces the computational workload, increasing the sensitivity of cross-linking site detection at single amino acid resolution (Bae *et al*, 2020, 2021).

While RBDome and RBPome methods have generated a wealth of valuable data, each has its own caveats and noise levels. Thus, there is a possibility of recovering many false positive hits (Bogdanow *et al*, 2016; Nesvizhskii *et al*, 2006; Bae *et al*, 2020). For example, although RBDome methods promise single amino acid resolution of binding site identification, there is a degree of uncertainty when it comes to mapping the cross-linked amino acid (Bae *et al*, 2020; Kim & Pevzner, 2014; Edwards, 2013). Moreover, a recent study has shown that UV cross-linked amino acids detected by these methods can also be indirectly cross-linked to RNA (Knörlein *et al*, 2022). Evidently, experimental validation of the findings is critical; however, the available methodologies are generally low throughput, making it challenging to quantify what fraction of RBDome data are biologically meaningful. An alternative approach would be to enhance the reliability of the experimental results using computional approaches. For example, one could calculate what fraction of cross-linked amino acids in RBDome data are in known RBDs (Queiroz *et al*, 2019; Bae *et al*, 2021, 2020) or interact with RNA in available crystal structures (Knörlein *et al*, 2022). To conduct a meaningful statistical analysis, however, a ground truth dataset is required that (ideally) consists of a large collection of high-resolution structures of protein-RNA complexes. However, such datasets are not readily available, especially for model organisms for which few protein-RNA complexes have been structurally characterised. This includes one of our favourite model organisms: *Staphylococcus aureus*. Furthermore, although extremely informative, ground truth datasets are not exhaustive, as they generally only contain relatively stable interactions that can be structurally characterised.

As an alternative, but also complementary, approach for assessing and enhancing the quality of experimental RBPome and RBDome data, we developed a Python computational pipeline (pyRBDome). This pipeline compares results from these high-throughput analyses against a large database of predicted RNA-binding residues. The pipeline generates this database for proteins of interest using a wide variety of different prediction tools that utilise distinct approaches for predicting RNA-binding sequences. Subsequently, the pipeline aggregates the results and putative RBSs are superimposed on (model) structures and other human-readable formats. When provided with RBPome data, the pipeline enables users to extract the most likely RNA-binders and identify amino acids most likely to bind RNA. When provided with a list of cross-linked peptides (RBD-Map, RBDome data), and amino acids (RBDome data), pyRBDome identifies the most common peptide motifs associated with RNA-binding and determines whether the data are significantly enriched for predicted RBSs by calculating 3D distances between experimental and predicted RBSs. By displaying Pfam domains (Mistry *et al*, 2021) identified in 3D structures, the user can easily determine the domains involved in the interactions. By clustering the cross-linking sites/peptides in domain structures, pyRBDome can identify interfaces within domains involved in RNA-binding. In conclusion, pyRBDome can reveal important mechanistic insights into RNA recognition, greatly facilitating further experimental validation of RNA-binding.

A second and equally important motivation for developing this pipeline was to make the analysis of RBP/RBDome datasets more accessible to groups that do not routinely perform such experiments or wish to analyse existing datasets. Moreover, because the pyRBDome code was written as Python Classes with associated test Jupyter notebooks, these can also be readily incorporated into new software tools.

Here we demonstrate how pyRBDome can effectively identify putative RNA-binding sequences in human and bacterial proteins and enhance RBDome datasets computationally. Moreover, using machine learning (ML), we show that combining prediction results from distinct computational tools employed in pyRBDome can enhance the sensitivity and specificity of computational prediction of RNA-binding amino acids in RBPs. We provide a detailed comparison with human structures of protein-RNA complexes, which revealed that UV cross-linking sites in proteins often correlate with the proximity to RNA in structurally characterised protein-RNA complexes, but not necessarily with direct RNA interaction.

## Results

### The pyRBDome pipeline

The main goal of this project was to develop a pipeline that would enable us to evaluate and enhance the quality of RBPome and RBDome datasets. The pyRBDome pipeline is written in Python, and the various analysis steps are provided in a series of Jupyter notebooks to facilitate the process of following, controlling and adjusting the analysis steps. The pipeline consists of two parts: pyRBDome-Core and pyRBDome-Notebooks. The former contains the Python classes and functions that are required for running the pyRBDome-Notebooks code. Each class in pyRBDome-Core has associated test Jupyter notebooks, making it easy to learn how to run the code. This should facilitate incorporation of the code into new bioinformatics tools. All the notebooks can be run either in Jupyter, or in the terminal using papermill (https://papermill.readthedocs.io/en/latest/). A schematic representation of the entire pipeline is shown in Fig. EV1. A minimum requirement for running the pipeline is a CSV file with a list of UniProt IDs for their proteins of interest. The pipeline will then enable users to identify putative RNA-binding amino acids within these proteins. If a list of putative RNA-binding peptides or amino acids for these UniProt IDs was provided, such as data from RBDMap (Castello *et al*, 2016), or RBS-ID (Bae *et al*, 2020, 2021), the pipeline will enable the user to identify which among the provided sequences/amino acids contains predicted RNA-binding residues, enabling effective selection of sequences that are likely to bind RNA. An example of such a CSV input file is provided in Dataset EV1. To facilitate these analyses, pyRBDome relies on multiple distinct RBS prediction tools. Considering the large size of RBS-ID and RBDMap data, and therefore the need to process a substantial number of proteins within a reasonable timeframe, the selection of these tools was based not only on their performance, but also on their runtime, and the ability to submit many proteins to webservers (also see Discussion).

RBS predictions are generally based on a wide range of features, such as amino acid sequence, structural data, and physicochemical properties of the studied proteins. Two of the computational programs used were specifically designed to identify potential RBSs using protein structure (aaRNA (Li *et al*, 2014)) and/or sequence information (aaRNA and RNABindRPlus (Walia *et al*, 2014)). However, a potential limitation of using these programs is that they were trained on data from known RNA-binding proteins (RBPs), which might make them less effective in identifying RNA-binding residues in unconventional RBPs. Therefore, we also analysed our data using BindUP, which predicts RBSs based on the electrostatic features on the protein surface and can more reliably detect non-canonical RBPs (Paz *et al*, 2016). RBSs can sometimes overlap with small molecule binding sites of enzymes, such as in the case of GAPDH, aconitase (Walden *et al*, 2006), and thymidine synthase (Chu *et al*, 1991). Hence, we used FTMap (Brenke *et al*, 2009) to find putative small molecule binding sites in structures. FTMap identifies possible ligand-binding pockets by globally docking a series of small organic probes onto the input structures to identify protein regions that represent binding hotspots. Incorporating FTMap data also offers the additional benefit of enabling the selection of RNA-binding proteins (RBPs) with a higher likelihood of being druggable. Additionally, many RBPs contain flexible and/or disordered domains, which are common in eukaryotic species. Therefore, we also included DisoRDPbind (Peng & Kurgan, 2015), which predicts RBSs in intrinsically disordered regions.

Consequently, pyRBDome integrates five independent yet complementary computational methodologies to compare against biochemically derived RNA-interacting protein sequences. While each approach has its own degree of uncertainty, our rationale lies in the consistency across these methods to identify amino acids more likely to be *bona fide* RBSs.

Several of the aforementioned tools rely on structural data to make their predictions. If available, the pipeline automatically downloads these structures from rcsb.org. In cases where such information is unavailable, pyRBDome retrieves structural estimates generated by AlphaFold2 (Jumper *et al*, 2021) or the homology modelling server SWISS-MODEL (Holm & Rosenström, 2010). This facilitates the analysis of RBPome and RBDome data from less well characterised model organisms.

To compare the experimental data to the predictions, for each peptide sequence provided, the pipeline calculates the minimal distance (in Å) to RBSs predicted by the individual tools. It stores its progress, such as whether files have been downloaded from webservers or specific tasks have been completed, as well as the analysis results in an SQLite database. The final results can subsequently be exported to CSV files where for each cross-linked peptide (Dataset EV2) or amino acid (Dataset EV3) provided, the pipeline reports where in the PDB file the peptide was mapped to and how frequently a predicted RNA-binding amino acid was detected. Manual inspection of the data in PyMOL revealed that cross-linked peptides and amino acids were often found near known RBSs. Therefore, we consider cross-linked sequences (peptides or amino acids) that are in close proximity of predicted sites (within hydrogen bonding distance (4.2Å) as a starting point) as promising hits. Thus, for each amino acid in each protein, the pipeline also reports its distance to predicted RBSs and distance to RNA molecules in known structures, if this information is available (Dataset EV5). Finally, using Interproscan (Quevillon *et al*, 2005), locations of domains within the protein sequences are determined, making it possible to identify domains involved in RNA-binding. The tables that are generated by the pipeline make it straightforward to statistically identify sequences obtained from RBDome experiments that are more likely to be *bona fide* RNA-binders.

### UV cross-linking data infrequently agrees with structural data

To showcase the feasibility of pyRBDome, we applied the pipeline to a recent human RBS-ID RBDome dataset (Bae *et al*, 2020). This dataset was chosen because, at the start of this project, it was the richest cross-linking dataset available: It includes data for almost 600 human RBPs and predicted RNA cross-linked amino acids for each protein. To facilitate the comparison of experimental data with predictions, pyRBDome requires peptide sequences that are at least 4 amino acids long as it needs to locate these sequences in 3D (model) structures. However, because the published RBS-ID data only provided the locations of cross-linked amino acids, we artificially extended these sequences on both ends with varying lengths (up to 27 amino acids; arbitrary number) to generate a dataset that we refer to as the “cross-linked peptide” dataset. The results of the pyRBDome analyses of this dataset is organised in tabular form in Dataset EV4.

If the user provides amino acid cross-linking data, the pipeline determines the preferentially cross-linked amino acids. Consistent with previous analyses (Bae *et al*, 2020), pyRBDome identified cysteines and the aromatic amino acids tyrosine, tryptophan, and phenylalanine as the most cross-linked amino acids (Fig. EV2A). Therefore, the user should expect to see a similar enrichment in their data. The pipeline performs the same analysis by grouping the amino acids into bins based on their physicochemical properties (Fig. EV2B), which identified sulphur-containing and aromatic amino acids as preferentially cross-linked. pyRBDome also enables the user to determine if sequences from specific domains were preferentially cross-linked. Using the InterProScan package (Jones *et al*, 2014; Blum *et al*, 2021) pyRBDome searches for domains within the proteins identified in the experimental data and it then counts how frequently cross-linked peptides and amino acids were mapped to these domains. Consistent with previous work (Bae *et al*, 2020), the canonical RNA recognition motif (RRM) and hnRNP K homology (KH) RBDs were the most enriched domains in the cross-linking data, followed by zinc finger (ZnF: C2H2, CCCH, and CCHC), WD40 repeats, and Helicase/DEAD domains (Fig. EV2C).

A second reason for choosing this human RBS-ID dataset was that high-resolution protein-RNA structures were available for 155 of the approximately 600 proteins. Consequently, we were able to compare the RBS-ID results with both RBS predictions collated by the pyRBDome pipeline and known protein-RNA interactions (ground truth dataset). Having ground truth datasets also allowed us to benchmark the different prediction tools employed in pyRBDome and to directly compare their performances (detailed below). To establish such human ground truth datasets, we downloaded hundreds of PDB files containing human protein-RNA complexes from rcsb.org. This yielded 371 protein-RNA structures (including the 155) that met our criteria for downstream analyses (see Methods for details). Using these structures, we generated two distinct ground truth datasets. Firstly, we used Protein-Ligand Interaction Profiler (PLIP; Adasme et al, 2021) to identify amino acids directly interacting with RNA in these structures. This ground truth dataset is referred to as GT-PLIP. The PLIP software package also enabled us to identify specific types of protein-RNA interactions, such as hydrogen-bonding, π-stacking, hydrophobic and salt-bridge interactions. However, due to limitations in resolution, not all structures generated PLIP results, yielding a relatively small dataset comprising of 192 proteins. To address this (potential) limitation, we established a second ground truth dataset, categorising amino acids that are within hydrogen-bonding distance (4.2Å) of RNA as RNA-binding (0 for non-interacting and 1 for interacting amino acids). We refer to this ground truth dataset as GT-Distance. This generated a richer and larger dataset (n=347), with ∼10% of the amino acids assigned as RNA-interacting. To capture all experimentally determined protein-RNA interactions for each protein, PLIP and distance-based detection of RNA-binding amino acids were performed using all available protein-RNA structures associated with individual UniProt IDs. Subsequently, the analysis results from multiple PDB files for a protein were merged into a single PDB file that stored for each amino acid the minimal distance to RNA and how frequently binding to RNA was detected.

To compare the performance of the prediction tools employed by pyRBDome, we used our ground truth datasets and recommended probability/scoring thresholds for identifying an amino acid as RNA-binding (Brenke *et al*, 2009; Li *et al*, 2014; Walia *et al*, 2014; Peng & Kurgan, 2015; Paz *et al*, 2016). The key performance metrics for each predictor (Fig. EV3). show that RNABindRPlus is one of the better performing tool on both the GT-PLIP and GT-Distance datasets, achieving the highest accuracy and precision. Notably, the performance of aaRNA on our GT-Distance dataset was comparable to its performance on a smaller ground truth dataset consisting of 67 RBPs (RB67; (Li *et al*, 2014)).

To simplify and automate the generation of ground truth datasets, we have included scripts in pyRBDome-Core that contain code needed for automated downloading of protein (FindUniProtPDBStructures.py) and protein-RNA complexes (FindUniProtRNPStructures.py) associated with specific UniProt IDs from rcsb.org, as well as code to calculate the distances of each amino acid to RNA (ProteinNAdistanceAnalyses.py).

We also wrote code to automate the PLIP analysis and the processing of the analysis results (https://git.ecdf.ed.ac.uk/sgrannem/pyDRBPNA). All the results generated by our ground truth analysis code is summarised in Dataset EV5. Illustrative examples of the ground truth datasets are showcased in Fig. 1A and 1B, presenting the outcomes within the crystal structures of the human SRP19 protein complexed with SRP RNA fragments.

**Figure 1.**
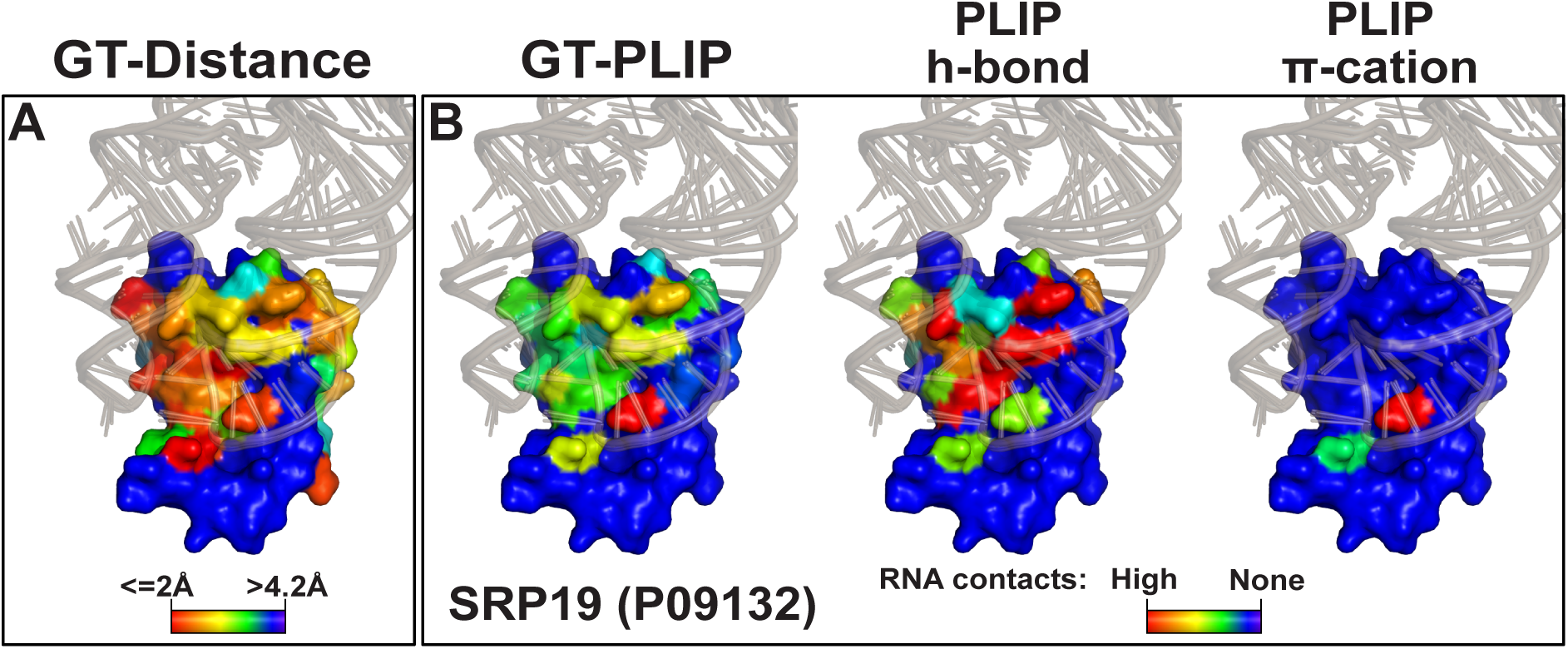
Ground truth analysis results for the human SRP19 protein. Shown is a surface representation of the structure of the human SRP19 protein in complex with a variety of co-crystallised RNA structures (wheat colour), obtained from available SRP19 protein-RNA complexes and superimposed on the protein structure. **(A)** Colouring amino acids in SRP19 by distance to RNA. Blue colours indicate amino acid residues more than 4.2Å away from RNA. The more the colour of the red spectrum, the closer the amino acid is to co-crystallised RNA in 3D. **(B)** As in (A) but colouring by how frequent an amino acid was detected to interact with RNA by PLIP in available structures.

To streamline the interpretation of the results, after the completion of the analyses, the pipeline generates PDB files that visually represent the prediction outcomes on the structural data, alongside PDF files containing the aligned prediction results within the protein sequence. It also generates convenient PyMOL session files making it easy for the user to visualise all the relevant PDB files simultaneously. The results for the SRP19 protein are shown in Fig. 2. Data for all the analysed proteins are available from our GitLab repository (https://git.ecdf.ed.ac.uk/sgrannem/). We have also included code in the pipeline that uses the InterProScan package (Jones *et al*, 2014; Blum *et al*, 2021) to search for domains within the proteins. If detected, the domains are highlighted in PDB and prediction outcome PDF files (Fig. 2B). The residue highlighted in yellow in Fig. 2B indicates the SRP19 amino acid cross-linked to RNA in the RBS-ID data.

**Figure 2.**
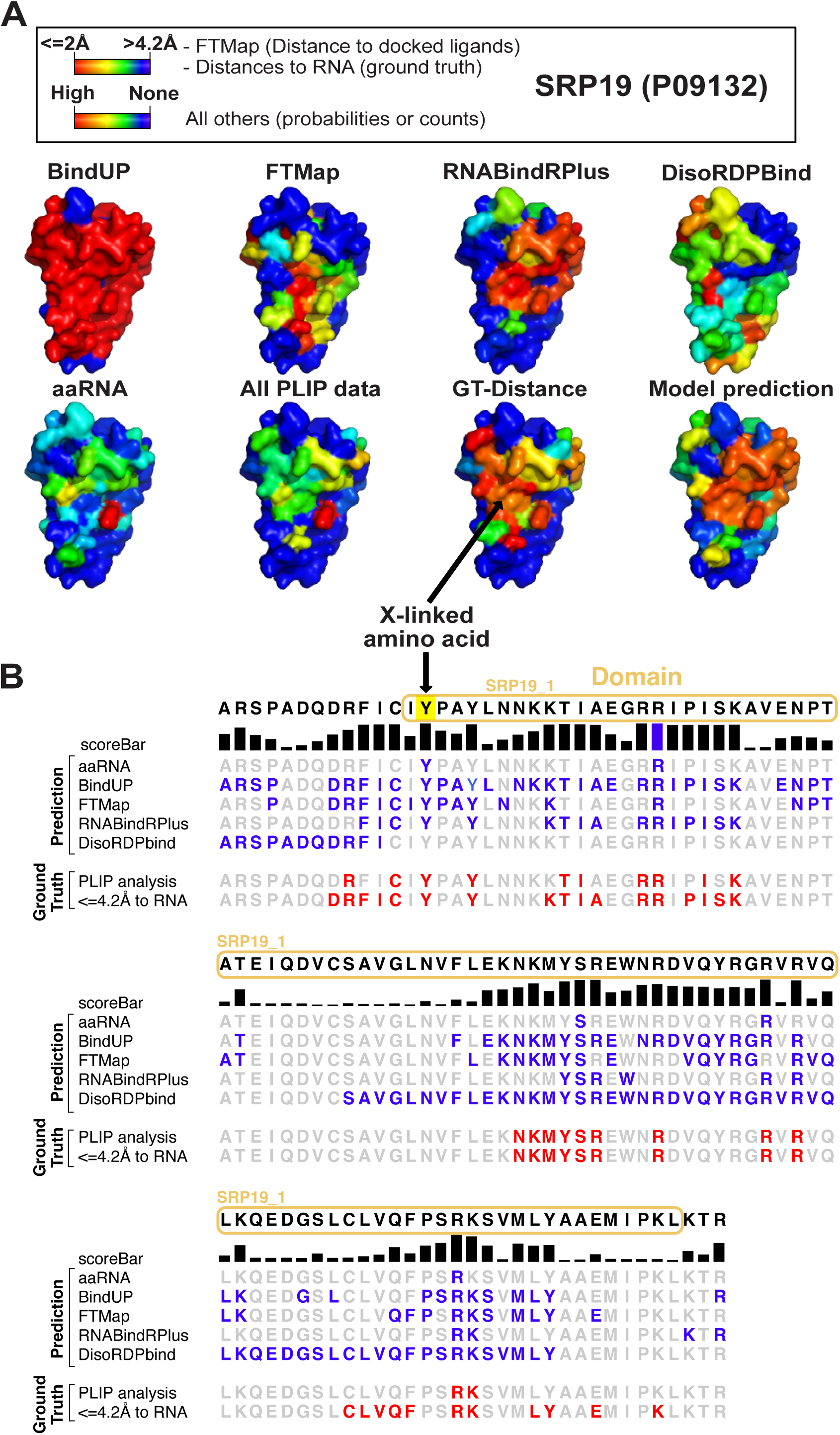
A representative example of pyRBDome analysis results. **(A)** Surface representations of the structure of the human SRP19 protein. Colours on the amino acids of SRP19 correspond to the scores/probabilities reported by different prediction algorithms. Blue colours denote amino acid residues with low scores, and the more the colour of the amino acid moves towards the red spectrum, the higher the RNA-binding probability/score. In the case of the FTMap results, the red-coloured amino acids are those less than 4.2Å away from docked small molecules, while blue colours indicate residues >4.2Å away from docked ligands. **(B)** An example of a pyRBDome PDF output file displaying the results along the linear sequence. Domains identified in the protein are outlined with ovals. Cross-linked amino acid residues are highlighted in yellow. The score bar represents the RNA-binding probabilities for the amino acid residues as determined by our XGBoost model using all the prediction results. The additional rows show results from various predictors (aaRNA, BindUP, FTMap, RNABindRPlus, and DisoRDPbind). Here, the blue amino acid residues indicate those with values at or above the recommended probability/score threshold (aaRNA: ≥0.18, BindUP: ≥10, RNABindRPlus: ≥0.5, DisoRDPbind: ≥ 0.16; FTMap <=4.2Å). The ground truth analyses results for SRP19 are also presented. GT-PLIP: red-coloured residues bind RNA in the SRP19-RNA structures. GT-Distance: red-coloured residues are amino acids positioned within 4.2Å of RNA in available structures.

### Aggregating data from multiple predictors increases confidence in RBS identification

The pyRBDome data analysis pipeline was founded on the principle that integrating outcomes from various distinct predictors not only enhances the quality of RBDome data but also enables more reliable identification of RBSs in proteins for which cross-linking data is absent. These assumptions were tested using machine learning (ML). Using the ground truth datasets outlined above, we developed eXtreme Gradient Boosting (XGBoost) ensemble classification models (Chen & Guestrin, 2016) that utilise the prediction results from the diverse tools used by pyRBDome as features to predict how likely an amino acid is to bind RNA (detailed in Fig. EV4). The XGBoost probability scores for SRP19, derived from all the pyRBDome results for this protein, are shown in the model prediction structure Fig. 2A and the score bar in Fig. 2B.

Developing a robust ML model for predicting RBSs is challenging, requiring extensive benchmarking against existing tools and deeply curated ground truth datasets, which is beyond the scope of this manuscript. However, precision-recall analyses (Fig. 3B and E) indicated that the XGBoost classifiers trained on the combined prediction results of the human ground truth datasets exhibited lower false positive and false negative rates compared to classifiers trained solely on data from individual tools. Furthermore, XGBoost models trained with more RBS prediction data displayed improved Area Under the Curve (AUC) values (Fig. 3C and F), implying they better distinguish between amino acids that bind RNA and those that do not. We note that models trained on GT-PLIP generally performed poorer than might be expected. This is likely because not all available structures could be analysed by PLIP due to limited resolution, reducing the size of the training dataset. Additionally, the unbalanced nature of GT-PLIP dataset, with only approximately 5% of all amino acids interacting with RNA, likely also significantly contributed to the lower precision of the XGBoost models trained on the PLIP data, despite artificially balancing the datasets (see Materials and Methods).

**Figure 3.**
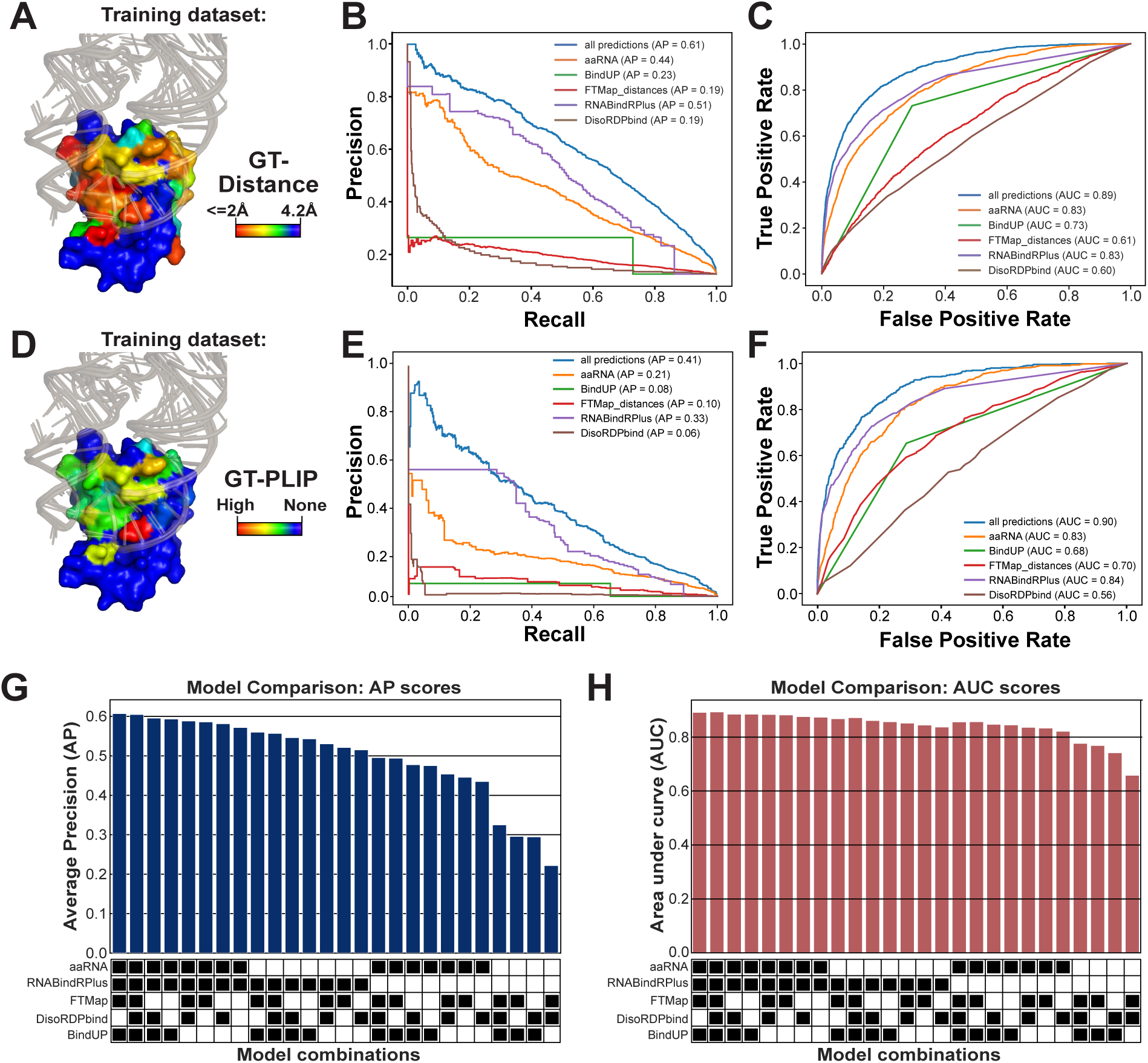
Assessment of XGBoost models trained on prediction models. **(A)** GT-Distance ground truth analysis results for the human SRP19 protein illustrating the distance in Å for each amino acid relative to RNA molecules. Shown is a surface representation of the structure of the human SRP19 protein in complex with a variety of co-crystallised RNA structures (wheat colour), obtained from available SRP19 protein-RNA complexes and superimposed on the protein structure. (colour gradient: red indicates a distance ≤2Å, yellow to green indicates a distance >= 2Å but < 4.2Å). **(B)** Precision-recall curves for the various XGBoost prediction models trained on the GT-Distance ground truth data using the predictions from either the individual tools or all predictions combined. The Average Precision (AP) score for each model is indicated in the legend (e.g., aaRNA AP = 0.46). **(C)** Receiver operating characteristic (ROC) curves for the same prediction models, with Area Under Curve (AUC) scores provided in the legend. **(D)** Visualisation of protein-RNA interaction predictions using an example from the GT-PLIP ground truth dataset, with the number of interactions identified by PLIP in available structures indicated in different colours (blue: none; green; at least 1, yellow, intermediate; red highest number). (**E-F**) Precision-recall (E) and ROC (F) curves for XGBoost models trained on the GT-PLIP ground truth data using predictions from the individual tools or all combined, with AP and AUC scores for each model shown in the legend. (**G-H**) Bar graph comparing the AP (G) and AUC (H) scores across different XGBoost models for the GT-Distance training dataset. The XGBoost models were trained using a combination of results from different predictors. The heat map below the bar plot indicates what predictions were used for training and testing the model.

It is important to note that the individual prediction tools (i.e., the model features) do not contribute equally to the predictions made by the XGBoost models, but the significance of each model is evaluated during the training. Analysis of the feature reliance in the performance of the XGBoost model (Fig. EV5A) revealed that BindUP, RNABindRPlus and aaRNA exhibited the highest importance among the RBS prediction tools, enabling the model to approximate the ground truth more accurately. Training XGBoost models using various combinations of RBS prediction data revealed that models trained with a more extensive collection of RBS prediction data showed increased precision (Fig. 3G; Average Precision (AP)). Notably, the AUC scores displayed less reliance on the number and type of RBS prediction datasets used.

These results validate our premise that combining results from multiple tools can improve prediction of RNA-binding amino acids in proteins and establish a strong foundation for the development of more enhanced ML models (see Discussion). These results also highlight the flexibility of our XGBoost model: even if the user is unable to provide results from some of the tools, the model will still be able to generate predictions with a reasonable average precision (Fig. 3G). We subsequently used the XGBoost model trained on the GT-Distance data to predict RBSs in proteins from the RBD-ID data. All the results from these analyses are provided together with the cross-linking information for each protein in Dataset EV4. On our GitLab repository we also provide PDB and PDF files summarising our XGBoost prediction results for all the proteins analysed during the course of the project.

### UV irradiation favours cross-linking RNA to positively charged and aromatic amino acids flanked by aliphatic residues

The likelihood of an RNA-protein interaction at a specific site is significantly influenced not only by the chemical properties of amino acids but also by its neighbours, owing to favourable protein folding or surface electrostatic forces. Recent studies have demonstrated that RBPs are enriched for tripeptide motifs consisting of positively charged, negatively charged, and aliphatic amino acids, and these triplets are conserved across evolution (Beckmann *et al*, 2015; Bressin *et al*, 2019). In three organisms that were analysed (*Homo sapiens*, *Escherichia coli* and *Salmonella. typhimurium*), tripeptides with a combination of arginines, lysines and glycines were strong predictors for RBPs. The pyRBDome pipeline can perform tripeptide motif analyses RBDome data, enabling users to identify motifs most likely to contribute to RNA-binding in their model organism. pyRBDome searches for tripeptide motifs enriched in the cross-linked peptides relative to randomly selected peptides from the same protein sequence (Fig. 4A). To enhance these analyses, pyRBDome also performs the same motif analyses based on the biochemical properties of the amino acids in the tripeptide motifs (Fig. 4C). Strikingly, the result show that while amino acids with positively charged residues are highly enriched in the human ground truth data (Fig. 4A, C), tripeptides containing combinations of aromatic (i.e., Y and F) and aliphatic (i.e., G, V and A) are very highly enriched in the cross-linked peptides (Fig. 4B, D). This is consistent with the strong bias towards UV cross-linking to specific amino acids, such as aromatic amino acids, to RNA.

**Figure 4.**
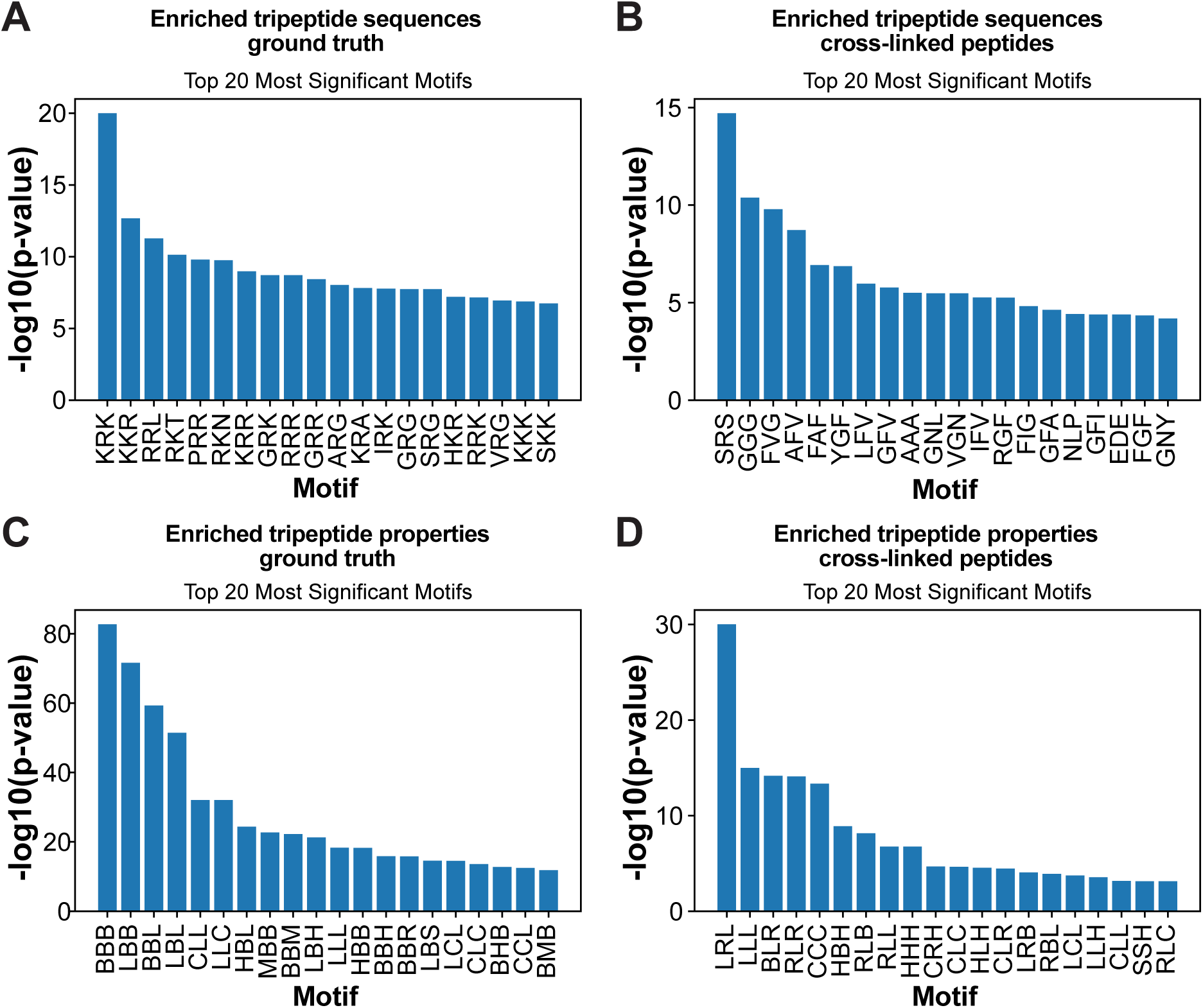
Cross-linked peptides are enriched for tripeptides containing aromatic and positively charged amino acids flanked by aliphatic residues. **(A)** Tripeptide motifs detected in RNA-binding regions (amino acids within 4.2Å from RNA) from known RBPs. **(B)** Tripeptide motifs enriched in the RBD-ID cross-linked peptides. **(C)** Enriched chemical properties of tripeptide sequences detected in the ground truth data described in (A). **(D)** as in (B) but now showing the chemical properties. Categories: L: aliphatic; R: aromatic; C: acidic; B: basic; H: hydroxilic; S: sulphur-containing; M: amidic. P-values were calculated using the Fisher exact test and corrected for multiple testing using the Benjamini-Hochberg procedure.

### pyRBDome reveals insights into domain RNA-binding interfaces

In addition to providing information about enriched domains in RBDome data, the pipeline can also identify RNA-binding interfaces within individual domains. UV cross-linking is inefficient and stochastic, so within individual protein domains, only a few of all possible RNA-binding interactions will be detected, providing limited mechanistic insights into domain-RNA interactions.

However, it is reasonable to assume that these domains within different proteins will have defined modes of RNA recognition. Therefore, if peptides/amino acids reported in RBDome data indeed represent genuine RNA-binding events, aggregating the cross-linking data from proteins that share the same domains may provide valuable insights into preferred RNA-binding interfaces.

To test this hypothesis, we further analysed the cross-linking data for RRM-containing proteins. The RRM domains in which cross-linking was detected were structurally aligned using MM-align (Mukherjee & Zhang, 2009) and superimposed. For those RRM domains for which crystal structures were not available, AlphaFold2 structure models were used. Subsequently, the cross-linked peptides and amino acids were highlighted within the superimposed structures (Fig. 5A-B). Typical RRM domains consist of four anti-parallel β sheets stacked on top of two α helices (Fig. 5A). Our analyses revealed that many cross-linked amino acids clustered in the same regions of the RRMs and concentrated in the β sheets (Fig. 5B). This finding is consistent with the essential role of the RRM β sheets in RNA-binding (Maris *et al*, 2005). Moreover, aromatic amino acids from the first and third β sheet that are important for RNA-binding (Maris *et al*, 2005) frequently cross-linked to RNA (Fig. 5C, red bars). However, to obtain meaningful results, many cross-linking events within a specific domain are required. To illustrate this point, the same analyses on type 1 KH domain proteins (36 cross-links), which were also enriched in the RBD-ID data, did not reveal a convincing cross-linking pattern (Fig. EV6). Nevertheless, our work demonstrates the potential of using high-throughput UV cross-linking studies for studying protein-RNA interfaces.

**Figure 5:**
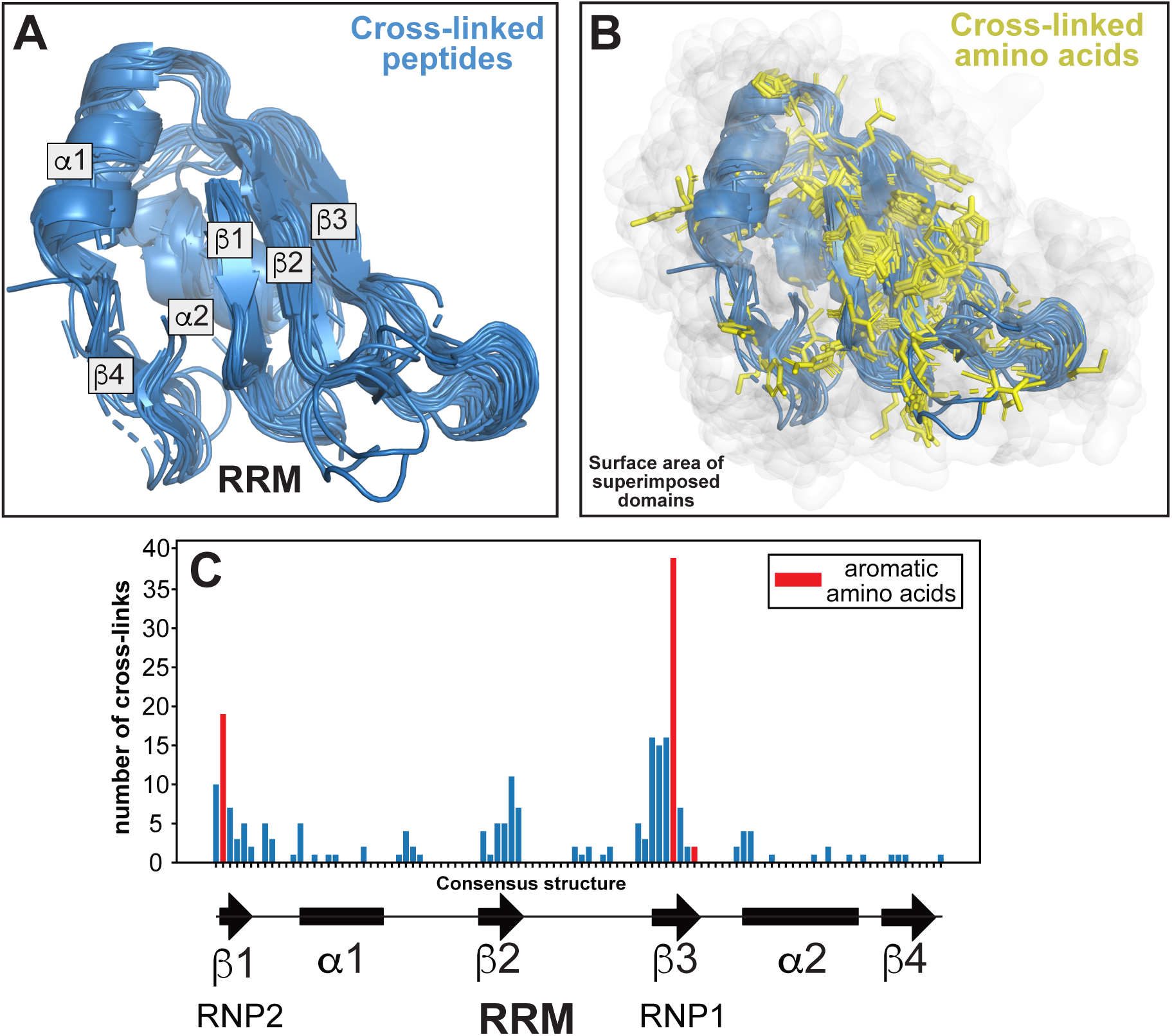
Insights into RNA-binding interfaces in protein domains through aggregated amino acid UV cross-linking data. **(A)** Superimposed peptide sequences mapped to RRM domains in proteins identified in the RBS-ID dataset. These sequences were aligned on available structural models of RRM domain-containing proteins. The various α and β secondary structural elements within the RRM domains are also indicated. **(B)** As in (A), but with the side chains of UV cross-linking sites within the domains highlighted as yellow sticks. The white cloud represents the surface area of the RRM domains. **(C)** The number of UV cross-links detected in all superimposed RRM domains (y-axis), correlating to their specific positions within the domain (x-axis). Below the x-axis, the consensus secondary structure for RRM domains is depicted for reference.

### UV-induced protein-RNA cross-links frequently occur in proximity to structurally determined protein-RNA contacts

We next asked to what extent the RBS-ID data agreed with our ground truth datasets. For this purpose, we only considered UniProt IDs from the RBS-ID data for which protein-RNA structures were available. We then compared this selection of RBS-ID data with our PLIP-analysed structures (GT-PLIP dataset). For each cross-linked amino acid reported in the RBS-ID data, we measured the distance (in Å) to the nearest RNA-binding amino acid detected by PLIP. The results were then aggregated into the cumulative plot shown in Fig 6A. Much to our surprise, these data showed that only 21.1% (43/204 amino acids) of the reported cross-linking sites interact with RNA in high-resolution structures (as reported by PLIP; Fig. 6A). Previous work (Knörlein *et al*, 2022) demonstrated that UV does not necessarily always cross-link the amino acids that in available structures bind RNA, but neighbouring amino acids can also be indirectly covalently attached to RNA. Consistent with this idea, more than half (56.4%) of the cross-linked amino acids were located within hydrogen-bonding distance (4.2Å) of PLIP sites and 42% within 4.2Å distance of RNA in these structures (Fig. 6B). Statistical analyses (Kolmogorov–Smirnov (KS) tests) revealed that RBD-ID data are indeed highly enriched for amino acid positions that are close to PLIP sites or RNA molecules in 3D structures (relative to shuffled cross-linked amino acids or all amino acids; Fig. 6A-B). These data therefore reinforce the idea that, when comparing the experimental data to existing structural data, UV cross-linking does not always capture amino acids directly binding to RNA, but that they are generally closer to RNA molecules.

**Figure 6:**
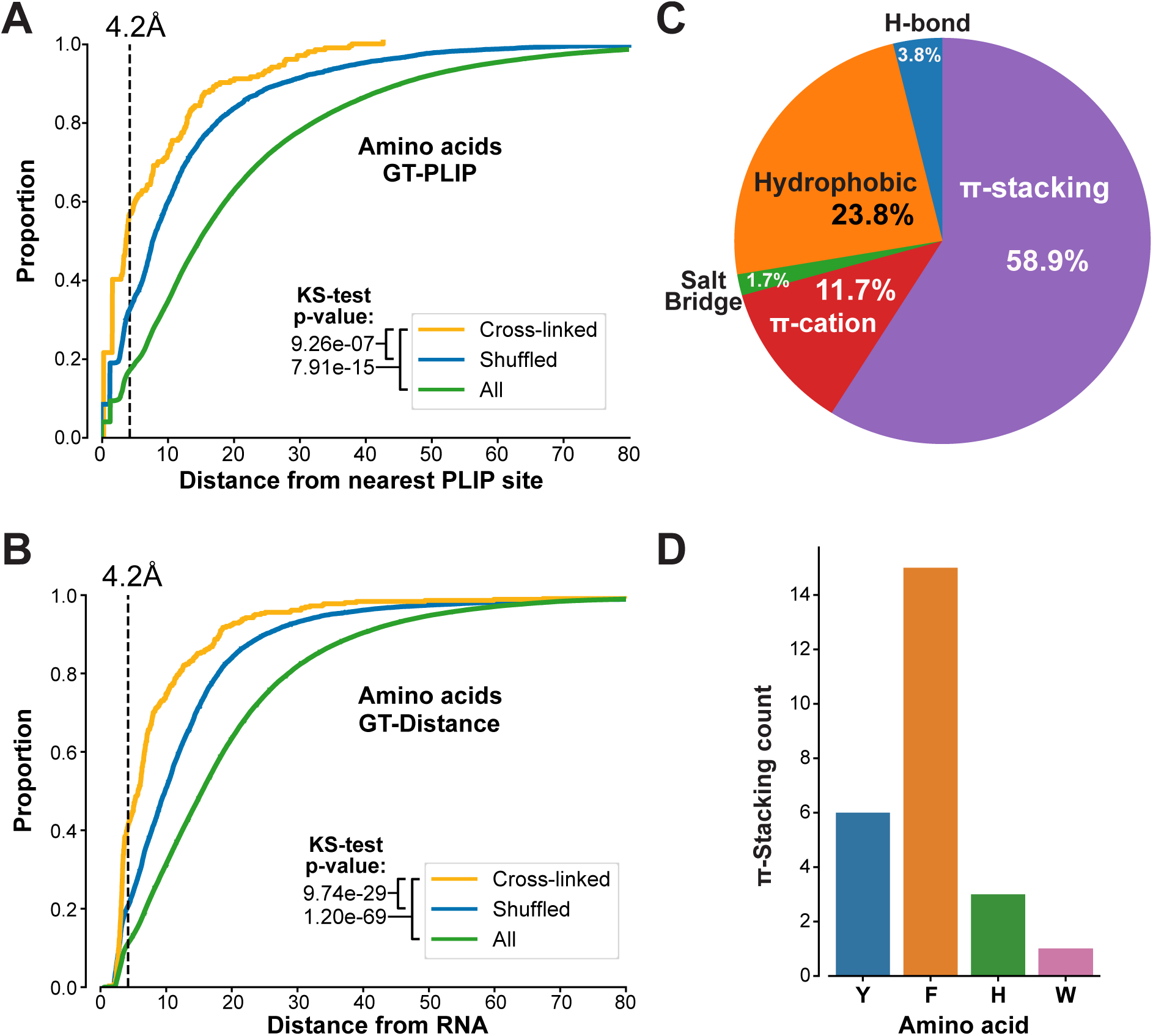
Limited concordance between UV cross-linking data and protein-RNA structures. **(A)** The cumulative distribution of distances for cross-linked amino acids (yellow), randomly shuffled amino acids (blue), and the total pool of amino acids (green), in comparison to established RNA-binding amino acids determined by PLIP. P-values, calculated using the Kolmogorov-Smirnov (KS) test, indicate significant differences between groups. The 4.2Å threshold, indicated by the dashed vertical line, is used to determine the proximity required for hydrogen bonding. **(B)** Similar to (A), this analysis plots the cumulative distances of cross-linked, randomly selected, and all amino acids within the studied RNA-binding proteins (RBPs), relative to their proximity to RNA. The KS test was also employed here to calculate p-values. **(C)** Amino acids that form π-stacking interactions are often cross-linked to RNA. The pie chart displays the percentages of each cross-linked amino acid involved in different types of interactions: hydrogen bonding (H-bond), π-stacking, π-cation, salt bridge, and hydrophobic interactions, as identified by PLIP. These percentages were calculated by dividing the number of a specific type of interaction by the total number of such interactions detected in the analysed structures. **(D)** Counts of cross-linked amino acids involved in p-stacking interactions. Y = Tyrosine, H = Histidine, F = Phenylalanine and W = Tryptophan.

We next focussed specifically on the cross-linked amino acids that overlapped with RBSs in our GT-PLIP dataset and asked what type of interactions they are involved in. Consistent with previous work (Knörlein *et al*, 2022), we find that phenylalanine π-stacking interactions with RNA are most abundantly detected (Fig. 6C-D). However, our results also suggest important contributions for hydrophobic and π-cation interactions (Fig. 6C).

### Cross-linked peptides as reliable proxies for RNA-binding regions?

As outlined above, a main reason why we established the pyRBDome pipeline was because for our model organism (Methicillin-resistant *Staphylococcus aureus*) there was an insufficient number of high-resolution structures of protein-RNA complexes available to generate a robust ground truth dataset for validation purposes. When analysing data from less well characterised organisms, the user can instruct the pipeline to determine whether cross-linked peptides and/or amino acids are highly enriched for RBSs predicted by the various tools employed by pyRBDome. Additionally, the user can test whether the cross-linking data is enriched for amino acids that, according to our XGBoost model, have high RNA-binding probabilities. Examples of such analyses on the human RBS-ID data are shown in Figures 7. These data indicate that the reported cross-linked amino acids have a significantly higher likelihood to bind RNA compared to randomly selected amino acids from the same proteins or the general population of all amino acids from the analysed proteins. However, the variability in the distribution of the RNA-binding probabilities for cross-linked RNAs, as shown by lower tail of the distribution, indicates that while cross-linked amino acids are indeed more likely to be predicted as RNA-binding, they are not a definitive indicator by itself.

**Figure 7:**
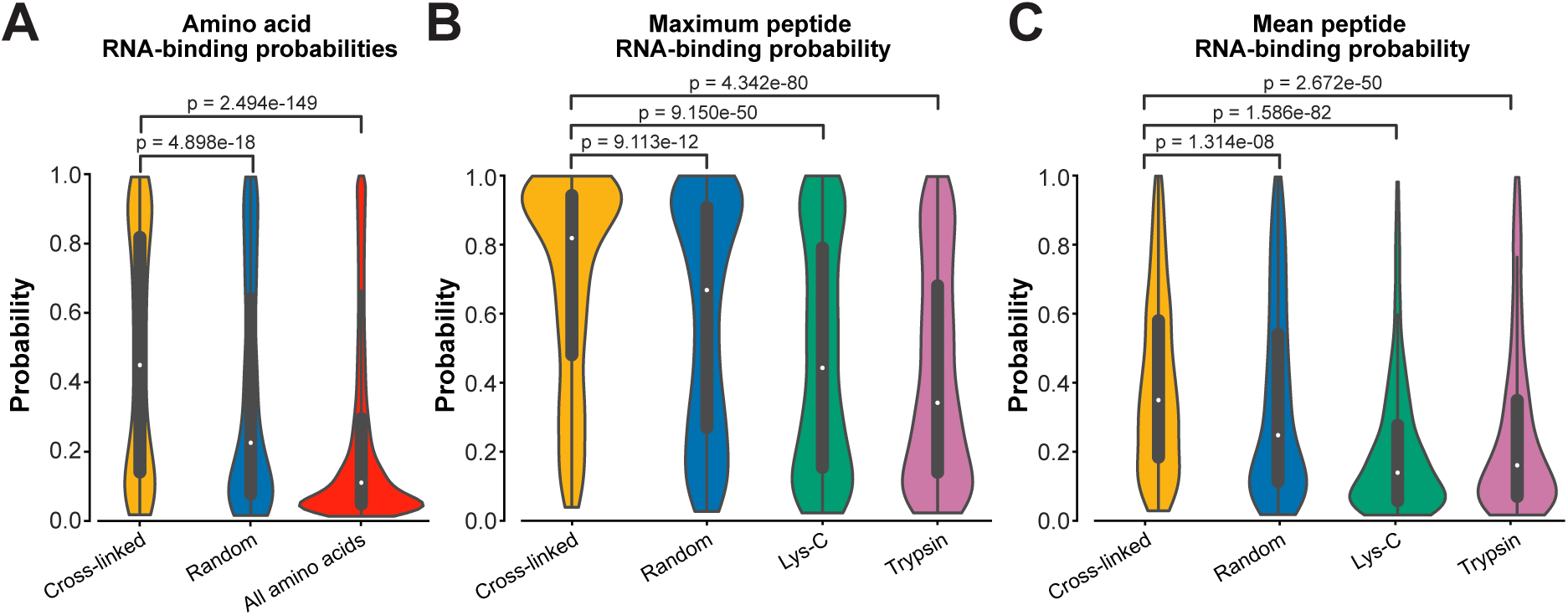
Cross-linked peptides as reliable proxies for RBSs. **(A)** Violin plots showing the distribution of RNA-binding probabilities as determined by our XGBoost model for cross-linked, randomly shuffled amino acids, and all available amino acids within the analysed RBPs. **(B)** The distribution of the highest RNA-binding probability score (determined by our XGBoost models) detected in cross-linked peptide sequences. Control datasets included randomly generated peptides with the same length distribution, and peptide libraries generated *in silico* by Lys-C or Trypsin digestion of the RBPs analysed here. **(C)** As in (B), but now for the average RNA-binding probabilities calculated for each cross-linked peptide. P-values, calculated using a two-sided Mann-Whitney-Wilcoxon test with Bonferroni correction, indicate significant differences between groups, as shown above each comparison. The violins represent density estimations of the distances, with wider sections indicating a higher frequency of amino acids at a particular distance. The white dot in the center of each violin plot denotes the median distance, and the thick lines within the violins represent the interquartile ranges.

Therefore, we next asked whether cross-linked *peptides* might be a better proxy for RBS detection. The pyRBDome pipeline allows the user to test this in two ways: Firstly, the user can compare the data with results obtained from individual predictors, such as aaRNA or FTMap, for example, as illustrated in Fig. EV7. These data show that the generated RBS-ID peptides were both enriched for predicted RBSs and/or more likely to be in closer proximity to these sites (aaRNA and RNABindRPlus; Fig. EV7A-B). Interestingly, the same was true for putative small-molecule binding sites predicted by FTMap (Fig. EV7C). The second approach determines whether the cross-linked peptides are enriched for amino acids with higher RNA-binding probabilities as determined by our XGBoost model. We addressed this by (I) tracking the highest RNA-binding probability found in a peptide sequence (Fig. 7B) and (II) calculating the mean RNA-binding probabilities for each peptide (Fig. 7C). Our analyses strongly indicate that cross-linked peptides typically include at least one amino acid with a significantly higher RNA-binding propensity compared to control samples (Fig. 7B). Notably, the RNA-binding probability distribution shown in Fig. 7B for cross-linked peptides is distinctly skewed towards higher values, suggesting that these peptides have a greater tendency for containing RNA-binding amino acids relative to the randomly selected control group peptides. However, the randomly generated peptides were not products of Trypsin and/or Lys-C digestion. To address this, we also compared the cross-linking data to peptides from parent proteins digested *in silico* by Trypsin/Lys-C. This comparison showed an even higher presence of predicted RBSs in cross-linked peptides, affirming the predictive strength of our XGBoost model and the significant value of cross-linked peptide data for detecting RBSs.

### pyRBDome correctly identifies RBSs in an *S. aureus* 3’-5’ exonuclease

Having extensively tested pyRBDome on human data, we next applied the pipeline on RBPome data from a less well characterised organism. For this purpose, we used our published RBPome data (Chu *et al*, 2022) generated on a clinically relevant *S. aureus* strain (USA300). (Model) structures for the top 200 enriched proteins were analysed by the pipeline and the results are available on our GitLab repository (https://git.ecdf.ed.ac.uk/sgrannem/pyRBDome_Notebooks_Staphylococcus_aureus_analyses). Given that our current XGBoost model had only been trained on human ground truth data, these analyses also tested the adaptability of the model to data from a genetically distant organism. To verify our findings, we focussed our analysis on the *S. aureus* polynucleotide phosphorylase (PNPase) 3’-5’ exonuclease, for which crystal structure data was available for both *S. aureus* (active site only) and *Caulobacter crescentus* (Hardwick *et al*, 2012; Wang *et al*, 2017). The latter structure also contained a short piece of RNA, enabling us to verify the reliability of the predictions.

To obtain a structure with the complete *S. aureus* PNPase sequence, we downloaded the AlphaFold2 model. This model was in good agreement with the published structures (RMSD values between 0.6 and 1; Fig. EV8A).

PNPase consists of three subunits that form a ring-like central channel where the RNA threads through the enzyme (Fig. EV8B). The S1 and KH domains, located at the C-terminus of each subunit, form the entrance of the channel, and direct single-stranded RNA towards the catalytic residues of the RNase PH-like domain, which is located at the N-terminal side of the channel (Hardwick *et al*, 2012). In the *C. crescentus* PNPase-RNA crystal structure, a 12-nucleotide RNA fragment interacts with the KH domain, through the conserved RNA-binding GSGG loop (Fig. EV8A-B, Fig. 8A-C). These amino acids were predicted to bind RNA with high probabilities by RNABindRPlus and our XGBoost model (Fig. 8A-C). The predictions of our pipeline largely accumulated on the internal surface of the ring-like structure that interacts with RNA. This can easily be observed when overlaying the RNA from the *C. crescentus* structure on the pyRBDome PNPase structure with the model predictions highlighted Fig. 8B-C. Interestingly, while FTMap highlighted the PNPase active site for its high potential to bind small molecules (Fig. 8A; red coloured amino acids), this region showed relatively low RNA-binding probabilities, reflecting the nuanced contribution of FTMap results to our XGBoost model’s predictions (Figs. 3 and EV5). The aaRNA analysis on the PNPase model structure did not yield any results and therefore these data were missing when using our XGBoost model, which was trained with aaRNA data, for predicting RBSs in this structure. Despite this, the XGBoost model yielded correct predictions for PNPase RNA-binding regions, again highlighting the degree of flexibility and robustness in the predictive capabilities of XGBoost models.

**Figure 8:**
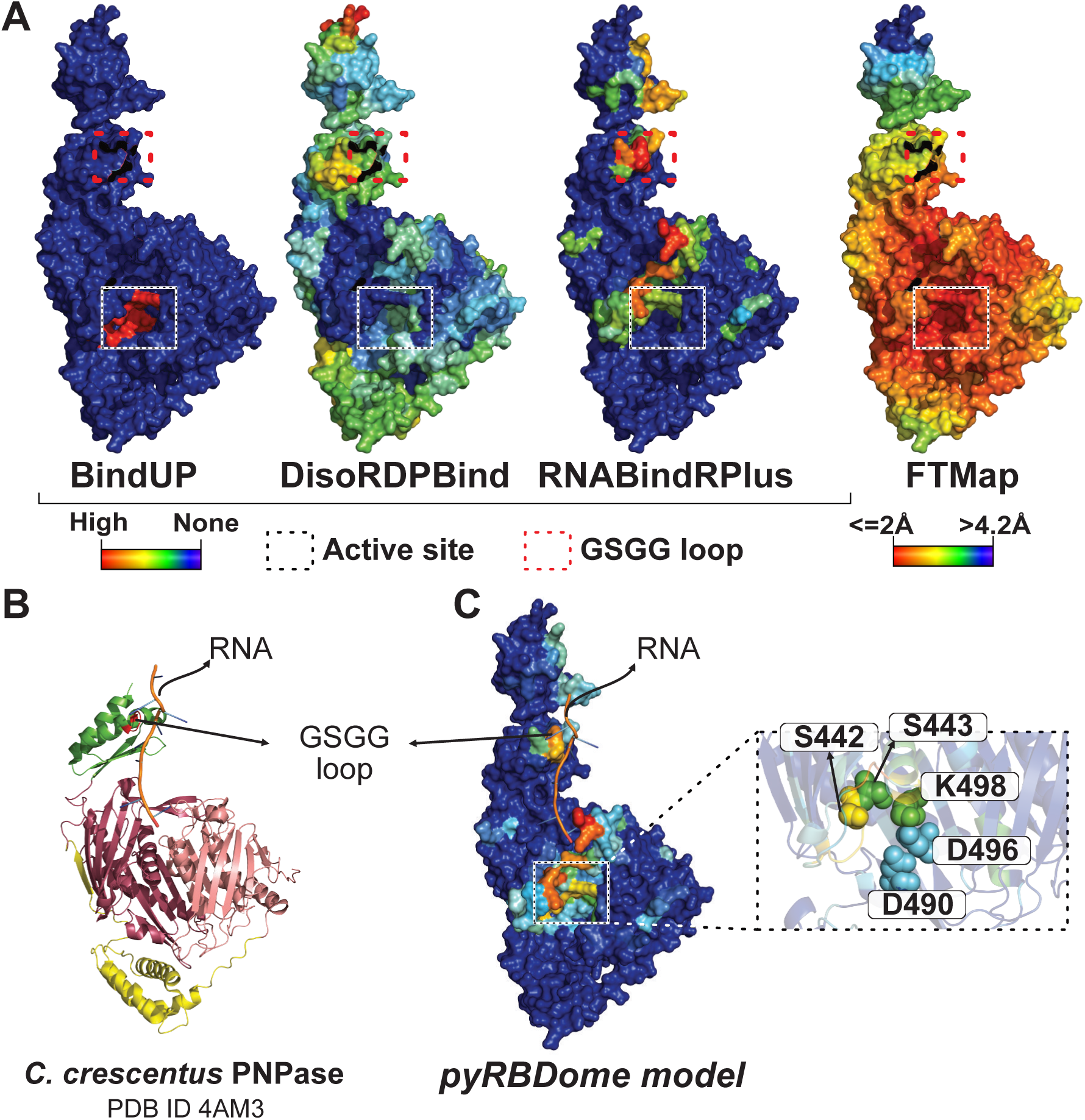
pyRBDome detects known RNA-binding regions in *S. aureus* Polynucleotide Phosphorylase (PNPase). **(A)** Results from prediction algorithms on the surface representation of a PNPase monomer. The colours for BindUP, DisoRDPbind, and RNABindRPlus results indicate RNA-binding probabilities, with cooler shades (blue) suggesting lower and warmer shades (red) indicating higher RNA-binding likelihood. For the FTMap results, warmer red shades signify shorter distances to docked molecules. The active site of the nuclease is marked with a square box. The GSGG loop is marked with a red square box. Blue colours represent amino acids with low RNA-binding prediction scores (BindUP, DisoRDPbind, or RNABindRPlus), whilst red colours indicate amino acids with high RNA-binding prediction scores. For the FTMap data, the blue to red colour gradient denotes decreasing distance to docked small molecules, with red indicating distances of ≤2Å and blue indicating distances of >4.2Å. **(B)** Crystal structure of PNPase from *C. crescentus*, in complex with RNA, PDB ID 4AM3 (Hardwick et al, 2012). The RNase PH-like domains, coloured in dark and light pink, are linked by a helical domain, coloured in yellow. The KH domain (green) interacts with the RNA of the structure through the GSGG loop (red). The S1 domain is absent from this crystal structure. **(C)** Structural alignment of the RNA from structure 4AM3 on the PNPase AlphaFold2 model with results from XGBoost model predictions trained on the prediction results from all algorithms. Catalytic residues are displayed as spheres and are highlighted in an enlarged view of the active site region.

In conclusion, the pyRBDome pipeline and the analysis tools we provide in this package are versatile and valuable tools for elucidating RNA-protein interactions across varied datasets and organisms.

## Discussion

Here we present the pyRBDome pipeline for *in silico* enhancement of RBPome and RBDome proteomics data. This pipeline, which leverages both protein sequences and structural information, employs a variety of distinct prediction tools for identifying putative RNA Binding Sites (RBSs) within target proteins (Fig. EV1 and EV4). It subsequently highlights the results from each prediction algorithm either within provided peptide/amino acid sequences, or entire protein sequences. The pipeline is capable of processing hundreds of proteins from large proteomics datasets or individual proteins. Significantly, the pipeline simplifies the complex data from these predictions, providing easily interpretable results that facilitate identification of residues involved in RNA-binding. The inclusion of PyMOL sessions allows users to visualise all the experimental and prediction results in 3D model structures simultaneously. Furthermore, pyRBDome includes statistical analyses to assess whether sequences obtained from RBDome studies show a significant enrichment of predicted RBSs, thus offering a quantitative measure that can improve the quality of the experimental data. Collectively, these findings underscore pyRBDome’s utility in streamlining the detection of RBSs in proteins and in effectively enhancing RBDome data.

### Agreement between RBDome UV cross-linking and structural data

To demonstrate the utility of pyRBDome we analysed a data rich human RBDome dataset (RBD-ID; (Bae *et al*, 2020), which provided, besides a list of (putative) RBPs, also an extensive list of RNA cross-linked amino acids. However, it did not contain the peptide sequences to which these cross-linked amino acids belonged. To address this, we artificially extended these amino acid sequences on both ends with varying lengths to create a peptide dataset suitable for analysis with our our pipeline. We found that both cross-linked peptides and amino acid sequences are significantly enriched in RBSs (RNA-binding sites), as predicted by individual tools or our combined XGBoost ensemble model. Surprisingly, when we compared the cross-linked amino acid data with our GT-PLIP dataset, which includes amino acids known to interact with RNA based on structural data, we observed limited overlap. While cross-linked amino acids were statistically more likely to be near RNA compared to randomly selected amino acids, only about 21% of them were actually found to bind RNA according to the available structural data. The limited overlap observed might suggest that UV cross-linking data contain a considerable amount of noise. However, it is important to note that our ground truth datasets, which were constructed solely from high-resolution structures, are also unlikely to include all possible protein-RNA contacts. Many structures contain proteins in complex with short pieces of RNA and therefore provide limited insights into the full RNA-binding capacity of the protein. Not every RNA substrate will also interact identically with an RBP and protein-RNA interactions can be highly dynamic and condition dependent. Though UV cross-linking can often capture such interactions *in vivo* and *in cellulo*, many of these might not be represented in static structures (also see (Bae *et al*, 2021)).

Our comparison of the RBS-ID data with our XGBoost model predictions, suggest that sequences of cross-linked peptides are more reliable indicators of RNA-binding sites than individual amino acids. This is because they tend to include amino acids with higher RNA-binding probabilities. Thus, comparing the cross-linking data with results from predictive models may offer a more effective solution for corroborating or supporting RBDome data. This is particularly true for models that are not solely reliant on existing protein-RNA structures for training. Such models are presumably better equipped to identify amino acids interacting with RNA, including those interactions not represented in structural data.

Another potential source of noise could stem from the analysis of mass spectrometry data. The software tools employed for analysing such datasets typically offer localisation scores, which indicate the probability of an amino acid being cross-linked to RNA. If the quality of a dataset is subpar, accurately pinpointing the precise cross-linking site becomes more challenging, leading to lower localisation scores and consequently, increased noise in the data. However, in the RBS-ID dataset that we analysed (Bae *et al*, 2020), 80% of the reported cross-linking sites (detected using MS-GF+ with a closed search; (Kim & Pevzner, 2014; Bae *et al*, 2020)) had very high localisation scores (between 0.8 and 1). While there is undoubtedly noise in the data, we would argue that the quality of this RBS-ID dataset is not a major contributor.

A recent study has also revealed that UV cross-linking does not exclusively target amino acids in direct contact with RNA; it can also affect those in indirect proximity (Knörlein *et al*, 2022). Furthermore, it was found that π-stacking interactions are key to directing the cross-linking reactions (Knörlein *et al*, 2022). This may also offer an explanation for our observation that few cross-linked amino acids were found to bind RNA in our GT-PLIP ground truth dataset, and if they did they were mostly involved in π-stacking. However, a significant proportion of the cross-linked amino acids were observed to be in close proximity to RNA within protein-RNA structures. Drawing on these findings and the bioinformatics analyses conducted in this study, when using pyRBDome data to design follow-up mutational analyses, we recommend prioritising aromatic, suphur containing and positively charged amino acids that have high RNA-binding prediction scores, that have undergone cross-linking or are located in cross-linked peptides, and those that are proximal to cross-linking sites, either sequentially or in the three-dimensional (model) structures.

### Developing an ensemble model for enhanced prediction RNA-binding amino acids

The foundational concept behind the creation of the pyRBDome pipeline stemmed from our belief that combining results from multiple predictors would improve the identification of RNA-binding residues in targeted proteins. While this was not the main goal of our project, the comprehensive datasets generated by pyRBDome presented a prime opportunity to validate this hypothesis through machine learning. By leveraging the predictive data from various tools, we developed a eXtreme Gradient Boosting (XGBoost) ensemble models. These models discern patterns within the aggregated predictive results and aligns them with known RNA-binding amino acids in the existing structural data. The main reasons for relying on XGBoost to build these preliminary models include its frequent outperformance of neural networks when presented with tabular data (such as the data used here), its ability to handle missing data points effectively (useful in cases where a protein could not be analysed by one of the prediction tools), its competence in dealing with unbalanced datasets (our ground truth datasets are unbalanced), and its tolerance to uninformative features (Chen & Guestrin, 2016; Grinsztajn *et al*, 2022). XGBoost therefore provided an excellent starting point for developing improved models for RBS prediction.

The preliminary models we constructed outperformed the individual tools, demonstrating greater accuracy and precision in predictions (Fig. 3). While these results are promising, there are areas where the XGBoost models could be further improved. For instance, our current models have exclusively been trained on data from human protein-RNA complexes. Therefore, their robustness could be enhanced by training the models on structurally characterised protein-RNA complexes (RNPs) from diverse organisms. It should also be noted that our training sets, in addition to AlphaFold2 models, mainly consists of structurally characterised proteins/domains. As a result RBPs with disordered RNA-binding regions are underrepresented or their disordered regions were excluded from the analyses. This underreprentation likely contributed to the less optimal performance of DisoRDPbind on our test data. However, this can be circumvented by reanalysing the data using *only* AlphaFold2 models, where these sequences will be represented (albeit not accurately folded). Alternatively, including a wider array of RNA-binding domains from disordered regions (Zhang *et al*, 2023) will undoubtedly enhance DisoRDPbind’s predictions and subsequently further improve the accuracy and precision of our XGBoost models. Therefore, the analyses presented here, constrained by the current datasets, do not fully capture the true potential of DisoRDPbind.

### Pipeline performance

The pyRBDome pipeline was designed to process a large number of proteins simultaneously, naturally leading to questions about the typical duration of an RBPome or RBDome dataset analysis. While there is no definitive answer, as it varies, performing the pyRBDome analysis on the RBS-ID dataset (consisting of 584 proteins) took approximately 8 days. The most time-consuming step involved submitting jobs to various servers, with tools like FTMap and aaRNA typically taking longer to yield results. The analysis duration primarily depends on factors such as the size of the proteins being analysed, the server’s computational power, and the server queue lengths. Despite these variables, we consider an 8-day turnaround to be quite reasonable for such a large dataset. Future developments of pyRBDome, as discussed in the next section, will focus on incorporating tools with shorter execution times. However, it’s important to note that faster processing does not always equate to more accurate results, presenting a constant trade-off.

### Future pyRBDome pipeline developments

To develop the pyRBDome pipeline, we evaluated a wide array of distinct tools designed to predict RNA-binding amino acids, and that take into consideration various sequence and structural features of ligand-binding proteins. However, integrating these tools into pyRBDome presented several challenges. These included inactive web servers and compatibility issues such as dependency conflicts and lack of comprehensive documentation, which hindered smooth integration with our Linux servers. Moreover, not all the web servers we tested were suitable for high-throughput analysis of protein sequences and structures, and some had run times that made the analysis of hundreds of proteins excessively time-consuming. This presented a notable challenge in integrating tools that could potentially outperform those currently described. However, the pipeline is continually evolving, and our existing Python code allows for relatively straightforward incorporation of new tools and the processing of their results.

Throughout this project, numerous advancements have been made in developing improved methods for predicting RNA-binding sites (RBSs) in proteins. A notable example is DeepDISOBind, an improved model for predicting RNA-binding residues in disordered regions (Zhang *et al*, 2022). We are in the process of incorporating the stand-alone version of this tool into pyRBDome-Core and pyRBDome-Notebooks. We are also testing PST-PRNA (Li & Liu, 2022), a deep learning model that predicts RBSs using protein surface topology. This tool outperforms aaRNA, a structure-based prediction method employed by pyRBDome. PST-PRNA has the added advantage of not relying on sequence identity and conservation for its predictions and may therefore perform better on non-classical RBPs. Preliminary data from these analyses can be found in versin 1.1.2 of our pyRBDome-Notebooks Ground truth analyses GitLab repository that details the development and analysis of our ground truth datasets (https://git.ecdf.ed.ac.uk/sgrannem/pyRBDome_Notebooks_Ground_truth_analyses). Other tools under evaluation are NCBRPred (Zhang *et al*, 2021), a sequence-based predictor likely to replace RNABindRPlus, and HybridRNAbind, a tool trained on both structural information and available RNA-binding regions in disordered domains (Zhang *et al*, 2023). We also tested HydRA (Jin *et al*, 2023), a deep learning method designed for detecting RNA-binding proteins and RNA-binding regions. Similar to the XGBoost model described here, HydRA functions as an ensemble classifier, utilising information from diverse prediction tools. It not only predicts a protein’s RNA-binding capacity, but can also detect potential RNA-binding regions in RBPs. Using our human GT-PLIP and GT-Distance ground truth datasets, HydRA’s performance in detecting RBSs was not as high compared to the individual tools employed by the pyRBDome pipeline or our XGBoost ensemble model (see 6.1.2_BinaryClassifierAnalysesRBDData.ipynb notebook in the pyRBDome-Notebooks Ground truth analyses repository). This is why we do not discuss the HydRA results here. This may be due to HydRA being optimised for predicting RNA-binding *regions*, whereas our ground truth datasets are more specific to individual RNA-binding *amino acids*. Despite this, we recognise HydRA’s value in identifying RNA-binding capacities in proteins and have incorporated code in version 0.2.0 of pyRBDome-Core and version 1.1 of pyRBDome-Notebooks to process and display HydRA predictions in PDF and PDB files. All the raw HydRa analysis results are also available on our pyRBDome-Notebooks GitLab repositories.

One might argue that constructing a pipeline dependent on multiple web servers, as in the case of pyRBDome, inherently invites reliability issues, as demonstrated by our experiences with inconsistent server availability. While our efforts are increasingly directed towards integrating standalone packages into the pyRBDome pipeline, it is important to acknowledge that running these prediction algorithms demands substantial computational resources. This includes the need for high-specification CPUs (Central Processing Units) and, more critically, GPUs (Graphics Processing Units). Not all research groups may have access to such computational facilities. Moreover, even for groups that do have such resources, the task of establishing and managing a pipeline comprising various stand-alone machine learning tools is very challenging as it involves dealing with numerous dependencies and configurations. Therefore, for future versions of the pyRBDome pipeline, we aim to strike a balance between utilising web servers and integrating standalone packages.

A longer-term goal is to make the results from analyses available in public databases with the aim to make the data more easily accessible for the wider public.

## Materials and Methods

### Repository content

A description of all the directories and type of files that the pyRBDome pipelines produce can be found in the README.md files in the individual repositories. The analyses described here used code from pyRBDome-Core version 0.2.0, pyRBDome-Notebooks version 1.0 and pyRBDome-Notebooks Ground truth analyses version 1.1.2.

### Generating the human ground truth dataset

We utilised the UniProt IDs from the RBS-ID dataset (Bae *et al*, 2020) to search rcsb.org for available protein-RNA structures. To expedite this process, we developed the script FindUniProtRNPStructure.py, which is now part of the pyRBDome-Core package. The code used for downloading these PDB files is available in the 1.0_FindRNPStructures_using_UniProt_IDs.ipynb notebook, located in the pyRBDome-Notebooks Ground truth analyses repository (https://git.ecdf.ed.ac.uk/sgrannem/pyRBDome_Notebooks_Ground_truth_analyses). For each UniProt ID, we retrieved protein-RNA structures that met specific criteria: a resolution of less or equal to 5Å and the presence of at least one RNA molecule. Owing to compatibility issues with CIF files, we chose to download only PDB files from rcsb.org. Each PDB file corresponding to a UniProt ID was then analysed to determine the minimum distance (in Å) of each amino acid to the RNA. We also developed a Python package that utilises the PLIP code (Adasme et al., 2021) to identify amino acids that interact directly with RNA in these structures. The code for conducting these analyses and a description of how to carry out such analyses is provided in the pyDRBPNA package on our repository. (https://git.ecdf.ed.ac.uk/sgrannem/pyDRBPNA).

To further refine these ground truth datasets, we merged the distance calculations and PLIP results for all PDB files associated with a single UniProt ID into a composite PDB file. This file records only the shortest distances to RNA for each amino acid in the b-factor column, as indicated in files ending with “distances_merged.pdb”. We also collated the frequency of RNA contacts by amino acids across the structures (as detected by PLIP), storing this information in the b-factor columns of files that end with “plip_merged_all.pdb”.

### pyRBDome package and pipeline description

The pipeline introduced in this paper consists of two parts: pyRBDome-Core (https://git.ecdf.ed.ac.uk/sgrannem/pyRBDome_Core) and pyRBDome-Notebooks (https://git.ecdf.ed.ac.uk/sgrannem/pyRBDome_Notebooks). The former contains all the scripts, functions, and classes that users need to execute the Jupyter notebooks. The code has been developed and tested extensively on Ubuntu Linux operating systems (OS) and can be adapted to work on Mac OS (12.7 and above). Details on how to install the packages and run the notebooks, and the required computational resources can be found in the README files on our repository. pyRBDome-Notebooks streamline the process of RNA-binding protein and cross-linking data analysis by automatically running predictions either online or locally. It then downloads, renames, and organises the results into specific directories. The pipeline stores any progress it has made as well as result from all the analyses in an SQLite database. This enables the user to keep track for which proteins (model) structures have been downloaded and whether these structures were analysed successfully by each prediction algorithm. Incorporating the SQLite database also enables the user to resume runs that may have failed or timed-out and helps avoid repeated submission of PDB files that have already been analysed. The results tables can also be easily exported to CSV files. All the notebooks can also be run sequentially in the terminal using papermill (https://papermill.readthedocs.io). Papermill is automatically installed when installing the pyRBDome-Core package.

The pyRBDome-Notebooks Jupyter notebooks each have their unique number. A detailed description of what analyses each notebook does is outlined below.

#### 1. Finding all available (model) structures for each UniProt ID

pyRBDome-Notebooks notebook 1.0_FindingPDBs.ipynb was used to download all available PDB files (<= 5Å resolution) associated with the UniProt IDs listed in the RBS-ID data (Bae *et al*, 2020) from rcsb.org (Berman *et al*, 2000), model structures that were generated by AlphaFold2 (Jumper *et al*, 2021) or the SWISS-MODEL webserver (Bienert *et al*, 2017; Guex *et al*, 2009; Studer *et al*, 2020; Waterhouse *et al*, 2018). For generating model structures, this notebook first queries the Alphfold2 database (https://alphafold.ebi.ac.uk) and downloads the latest model associated with that UniProt ID (PDB files ending with “_AF.pdb”). If it is unable to find any models, it submits the protein sequence to SWISS-MODEL. Only models with GMQE score higher than 0.7 were considered and their PDB files downloaded. Note that SWISS-MODELS were not used in this study. For proteins that could not be modelled by SWISS-MODEL or had a model of insufficient quality the protein sequences were blasted against the AlphaFold model organism genome (notebook 1.1_FindingPDBsViaSequence.ipynb) to identify the closest homologue (notebook 1.2_GetAlphaFoldModels.ipynb). In these cases, we only considered proteins that had a homolog with an identity of >= 99%. The PDB IDs associated with each protein are then saved in the available_PDBs table in an SQLite database (pyrbdome_full.db). The tables in the database have information about whether the PDB file was successfully downloaded and what chain is included in the PDB file.

#### 2. Getting protein domains from Pfam

After all the PDB files have been downloaded, notebook 1.3 will use the Interproscan tool (Jones *et al*, 2014; Blum *et al*, 2021) to download all the domain information associated with these proteins. Only Pfam domains are considered. A Linux version of Interproscan is provided in pyRBDome-Notebooks programs folder. The user will need to install a different version if Mac OS operating systems are used for the analyses.

#### 3. Creating peptide control datasets

Notebook 1.3 takes the protein sequence from each PDB file and digests the sequences *in silico* with Trypsin and Lys-C to generate a library of all possible peptides that could theoretically be detected by the mass-spectrometer for the protein of interest. If cross-linked peptide sequences were provided, notebook 1.4 will generate a library of random peptide sequences that are peptides of the exact same length distribution as the cross-linked peptides, but that were randomly extracted from the protein sequence.

#### 4. Performing RNA/ligand-binding sites predictions

To predict RNA/ligand-binding sites on the proteins of study, we chose five different prediction algorithms: aaRNA, BindUP, FTMap, RNABindRPlus and DisoRDPbind (Walia *et al*, 2014; Peng & Kurgan, 2015; Paz *et al*, 2016; Mehio *et al*, 2010). These notebooks will automatically submit all the PDB files to the respective web servers, download the results, and store the progress they have made with the analyses in the SQLite database. To further increase the performance of the pipeline, we are also implementing the PST-PRNA deep learning approach (Li & Liu, 2022) in our notebooks, which predicts putative RNA-binding amino acids entirely using the surface topology of the proteins in the structures. Preliminary results from these analyses are available in pyRBDome-Notebooks version 1.2.

#### 5. Mapping the cross-linked amino acid and peptide sequences to the PDB files

Notebook 3.0 takes the cross-linked, *in silico* digested and random peptide sequences and maps them to the PDB files. Once the peptides have been mapped, it will determine the location of cross-linked amino acids, if this information was provided. For example, if the peptide sequence “PSRKDPKYREWHHFL” is analysed by this notebook and it could be mapped to a PDB file sequence, it will record the start and end residue numbers for the peptide and what chain it was mapped to in the PDB file. For this example, the code returned the following result: 74A_psrkdpkyrewhhfl_88A. This shows that the peptide was mapped between residues 74 to 88 of chain A in the PDB file. Note that not all peptides will be mapped as many structures do not contain the complete protein sequence.

#### 6. Processing the results and storing them in PDB files

Notebook 4.0 collects all prediction results and any domain and mapped peptide/amino acid information and stores the results in the b-factor columns of the PDB file. This makes it possible to visualise the results in PyMOL or other viewers.

#### 7. Distance analyses

The series 5 notebooks take all the prediction results, map these to the peptide sequences and calculate the closest distance of the cross-linked peptides or control peptide sequences to amino acids predicted to be involved in RNA-binding. The results are stored in tables in the SQLite database. These tables enable the user to easily extract peptide sequences that contain predicated RNA-binding amino acids. For example, if it found a predicted RNA-binding amino acid in a mapped peptide (e.g. 74A_psrkdpkyrewhhfl_88A), it will indicate the location of this amino acid in upper case (e.g.74A_psrkdpky**R**ewhhfl_88A).

#### 8. Sanity check

Notebook 6.0 then looks at all the distance analyses and double checks if no errors were made in the calculations. This notebook is tremendously useful for troubleshooting any issues that might appear during the analyses.

#### 9. Analysis of cross-linked peptide and amino acid sequences

The series 7 notebooks search for enriched tripeptide motifs enriched in the cross-linked peptides and enriched amino acids in the cross-linked amino acid data, if available. It returns a table containing the sequences of the enriched amino acid motifs or chemical properties and associated p-values.

#### 10. Making the final output files

The series 8 notebooks gather all the prediction and cross-linking information from the PDB files that were produced by notebook 4 and place the information in a large table where RNA-binding probabilities provided by each algorithm are stored as well as the location of cross-linked peptides and amino acid residues. The notebooks in the pyRBDome-Notebooks analyses of the ground truth dataset also contain extra code that adds the distances to RNA molecules for each amino acid for all protein-RNA structures that were analysed. Notebooks 8.0 and 8.1 take all the prediction results available in the large table, feeds that to our XGBoost models, and calculates for each amino acid in each protein a probability for RNA-binding. The 8.2 statistical analysis notebook determines whether cross-linked peptides and amino acids (where available) are significantly enriched for predicted RBSs compared to the random peptide datasets and the peptides generated by Trypsin/Lys-C digestion of the protein sequences. Notebook 8.3 takes all the analysis results and produces a PDF file summarising all the results in the protein sequence for each protein. The scorebars in the PDF files indicate the XGBoost RNA-binding probabilities for each amino acid. Notebook 8.4 generates PyMOL session files that enables the user to conveniently load all PDB files into a single PyMOL session.

#### 11. Binary classification analyses. Training of XGBoost models

The ground truth pyRBDome-Notebooks ground truth analysis repository contains notebooks 6.1.1 and 6.1.2 outlining how the XGBoost models were trained on the GT-PLIP and GT-Distance ground truth datasets, These notebooks also include details about what parameter optimisation steps were performed and tests for analysing overfitting. The GT-PLIP and GT-Distance ground truth datasets are provided on our repository as a text file (https://git.ecdf.ed.ac.uk/sgrannem/pyRBDome_Notebooks_Ground_truth_analyses/-/blob/main/analysis_results/All_combined_results.txt) and the Datasets EV5 in the Supplementary Data. These files contain the names of the UniProt IDs that were analysed, the PDB files we used, a list of all the amino acids and residue numbers for ech protein in the PDB file, the distance of an amino acid to RNA (if available) and results from the PLIP analyses. Dataset EV4 also contains all the prediction scores from the individual tools for each amino acid.

For the training of the XGBoost ensemble model, we normalised the scores or probabilities from each individual predictor (aaRNA, RNAbindRPlus, BindUP, and DisoRDPbind) to a range between 0 and 1, where necessary. These normalised values were then utilised as feature values for training the models (Fig. EV4). In the case of FTMap data, the distances to docked molecules (in Å) were normalised to values between 0 and 1, with the highest values assigned to the shortest distances. The XGBoost model subsequently generates output files containing probabilities that indicate the likelihood of each amino acid interacting with RNA. Given that the number of RNA-interacting amino acids in the GT-PLIP and GT-Distance ground truth datasets was approximately 5-10%, we undersampled the majority class (i.e., non-interacting amino acids, labelled as ’0’s) in our training data to address the unbalanced nature of the dataset. To build the models, 80% of all structures in the ground truth datasets were used for training and 20% for testing. Utilising Python’s Scikit-learn and the Optuna optimisation framework (Akiba *et al*, 2019), we optimised the hyperparameter for our XGBoost models. This optimisation included 10-fold cross-validation to enhance the robustness and generalisability of the models. All models, including those trained on various combinations of prediction results, are available from our repository (pyRBDome-Notebooks Ground truth analyses; 6.1 series notebooks and folder ’xgboost_models’)..

#### 12. Analysis of predictions and cross-linking sites onto protein domains

Notebook 9.0 analyses (1) what domains were detected in cross-linked peptides and (2) which ones were enriched in the data. Notebook 9.1 extracts selected domains from the available PDB files, superimposes them and highlights prediction scores, cross-linked peptides, and cross-linked amino acids within the superimposed structures. To be able to run notebook 9.1, we added the Linux version of MMalign (Mukherjee & Zhang, 2009) to the ‘programs’ folder in the pyRBDome-Notebooks repository. This version was compiled on Ubuntu 22.04 and may not be compatible with later versions of Ubuntu and different operating systems. These analyses enable the user to determine whether predicted RBDs show specific cross-linking patterns, making it possible to gain information about domain RNA-binding interfaces.

## Data Availability

All the code and data analyses results are available from our GitLab repository (https://git.ecdf.ed.ac.uk/sgrannem) without restrictions. All the prediction and ground truth analysis results can be found on the repositories starting with pyRBDome-Notebooks. The pyRBDome-Core repository contains all the code required to run the pyRBDome-Notebooks Jupyter notebook files. The results of all the analyses are also available as Microsoft excel spreadsheets in the Supplementary information (Datasets EV2-5).

## Acknowledgements

We would like to thank Shaun Webb and Rasna Walia for help with RNABindRPlus, Songling Li and Daron Standley for help writing Python code for automatically submitting aaRNA jobs, Yun Zhou for helping with the analysis of PDB files, and Guido Sanguinetti, Andrea Weiße and Alfredo Castello for critically reading the manuscript. This work was supported by a Medical Research Council Non-Clinical Senior Research Fellowship (MR/R008205/1 to S.G.), a Darwin Trust award (N.C) and a Wellcome Trust PhD training fellowships awarded to H.M. (220406/Z/20/Z).

## Author contributions

**Liang-Cui Chu:** Conceptualisation; software; formal analysis; investigation; writing - original draft. **Niki Christopoulou:** Conceptualisation; software; formal analysis; investigation; writing - original draft; writing – review and editing. **Hugh McCaughan:** Conceptualisation; software; formal analysis; investigation; writing - original draft; writing – review and editing. **Sophie Winterbourne:** Conceptualisation; software; formal analysis; investigation; writing – review and editing. **Davide Cazzola:** Conceptualisation; formal analysis; investigation. **Shichao Wang:** Conceptualisation; software; formal analysis; investigation; visualisation; methodology. **Ulad Litvin:** formal analysis; investigation; visualisation. **Salomé Brunon:** formal analysis; investigation; visualisation. **Patrick Harker:** formal analysis; investigation; visualisation; writing – review. **Iain McNae:** Conceptualisation; investigation; methodology; writing – review; supervision. **Sander Granneman:** Conceptualisation; software; formal analysis; investigation; visualisation; methodology; writing – final draft, review and editing; supervision; funding acquisition; project administration.

