## Supplementary_information for "pyRBDome: A comprehensive computational platform for enhancing and interpreting RNA-binding proteome data"

### Expanded View Figures and Figure Legends

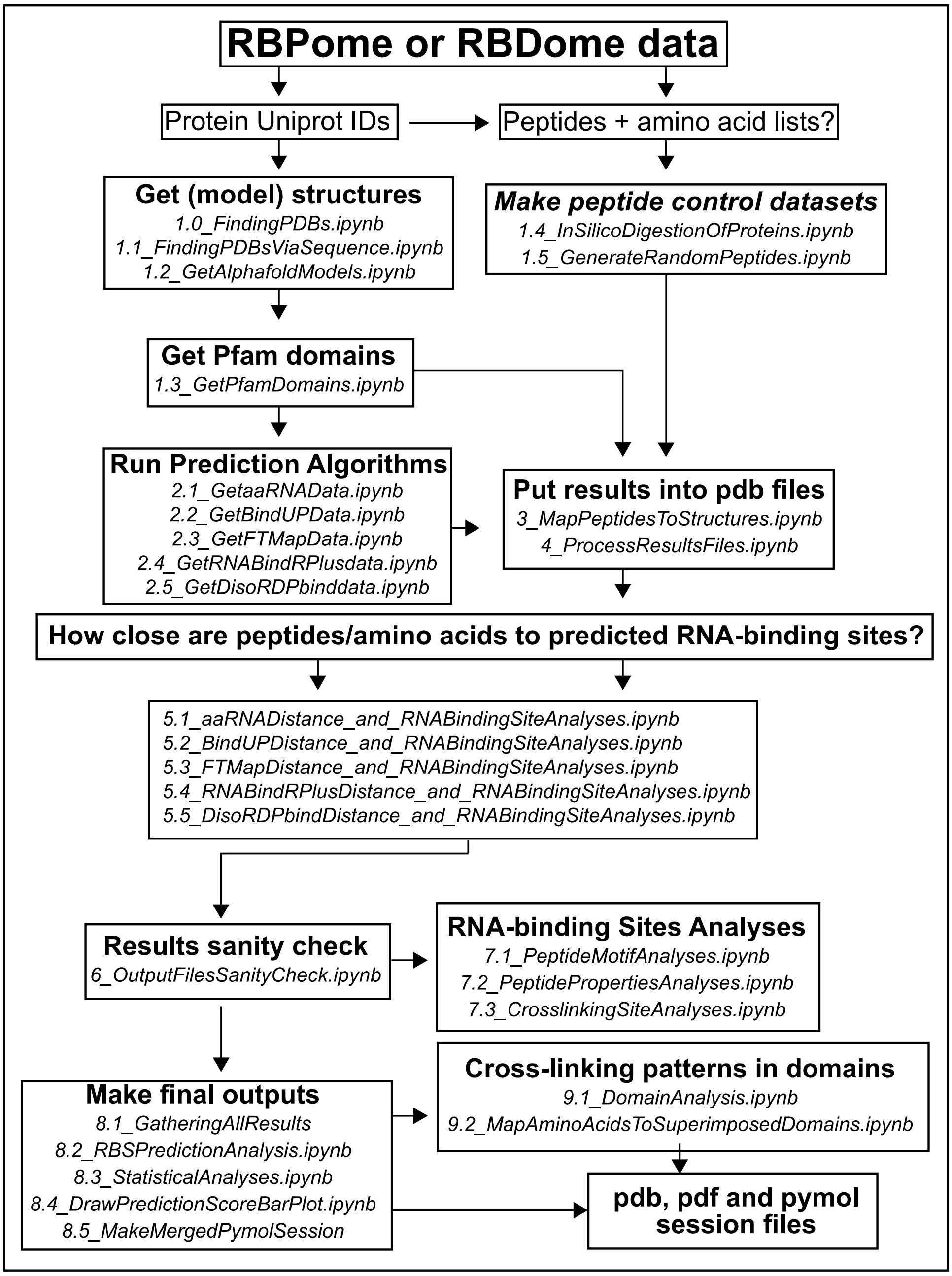


**Figure EV1: Schematic representation of the complete pyRBDome pipeline.** For a detailed description, please see the main text and the Methods section. Briefly, to start running the pipeline, a CSV file containing UniProt IDs is a minimum requirement. Information about cross-linked peptide and amino acid sequences can also be included. An example of an input file can be found on our git repository (https://git.ecdf.ed.ac.uk/sgrannem/pyRBDome_Notebooks/-/blob/main/pyRBDome_analyses/RBSID_human_data.xlsx). Each discrete analysis step in the pipeline is indicated with boxes. The names ending with. ipynb indicate the names of the Jupyter notebooks that are used in each step of the analysis.

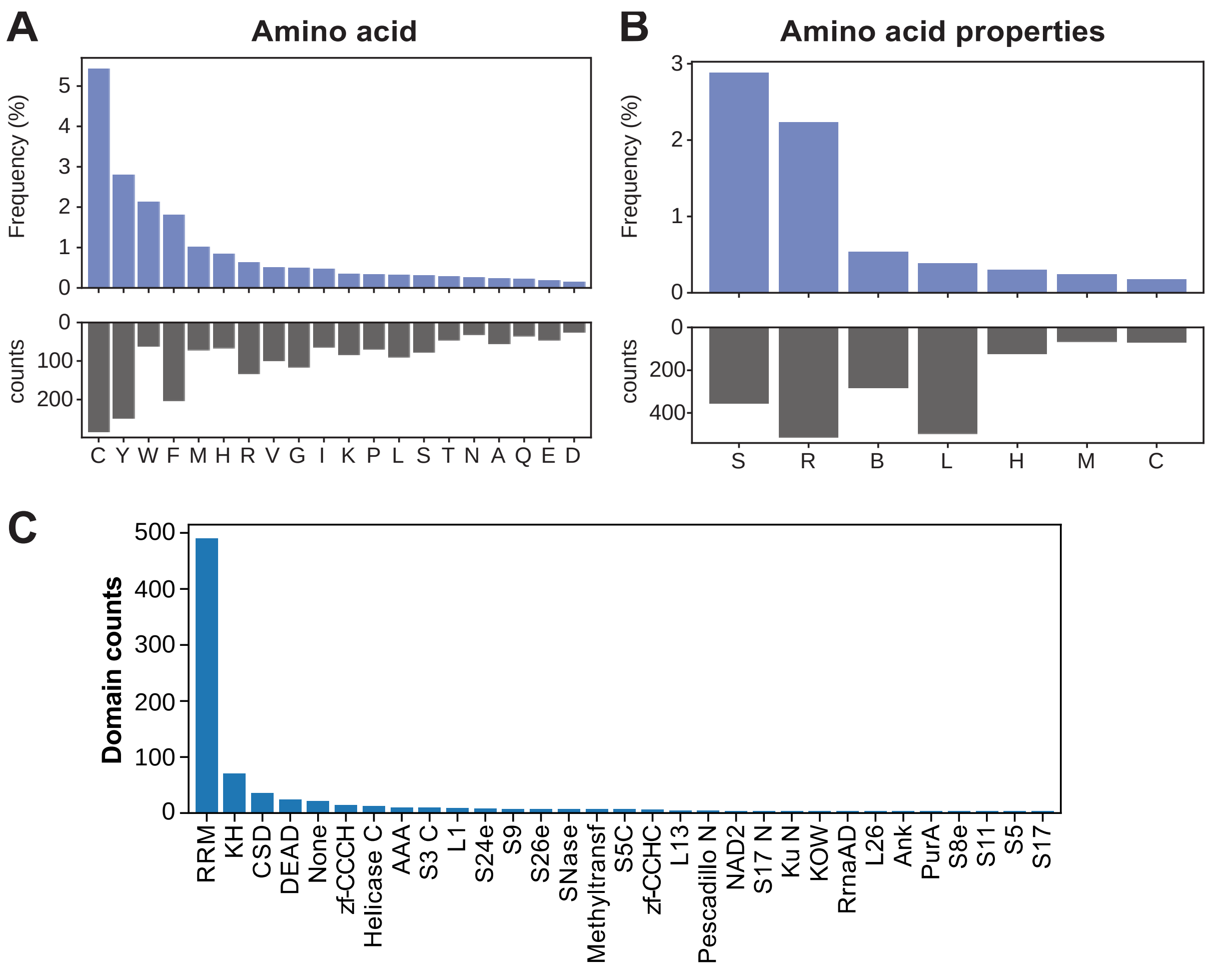


**Figure EV2: Analysis of amino acid and domain cross-linking preferences in RBD-ID data.**

(**A**) Counts (black bars) and frequency (blue bars) of cross-linked amino acids. Frequency is calculated by dividing the total counts of each amino acid observed in the cross-linking data by the total occurrence of that amino acid in the protein sequences of the analysed proteins. (**B**) Same as in (A) but now for the chemical properties of the amino acids. Categories: L: aliphatic; R: aromatic; C: acidic; B: basic; H: hydroxilic; S: sulphur-containing; M: amidic. (**C**) Histogram displaying the total number of times a cross-linking peptide was detected in specific protein domains.

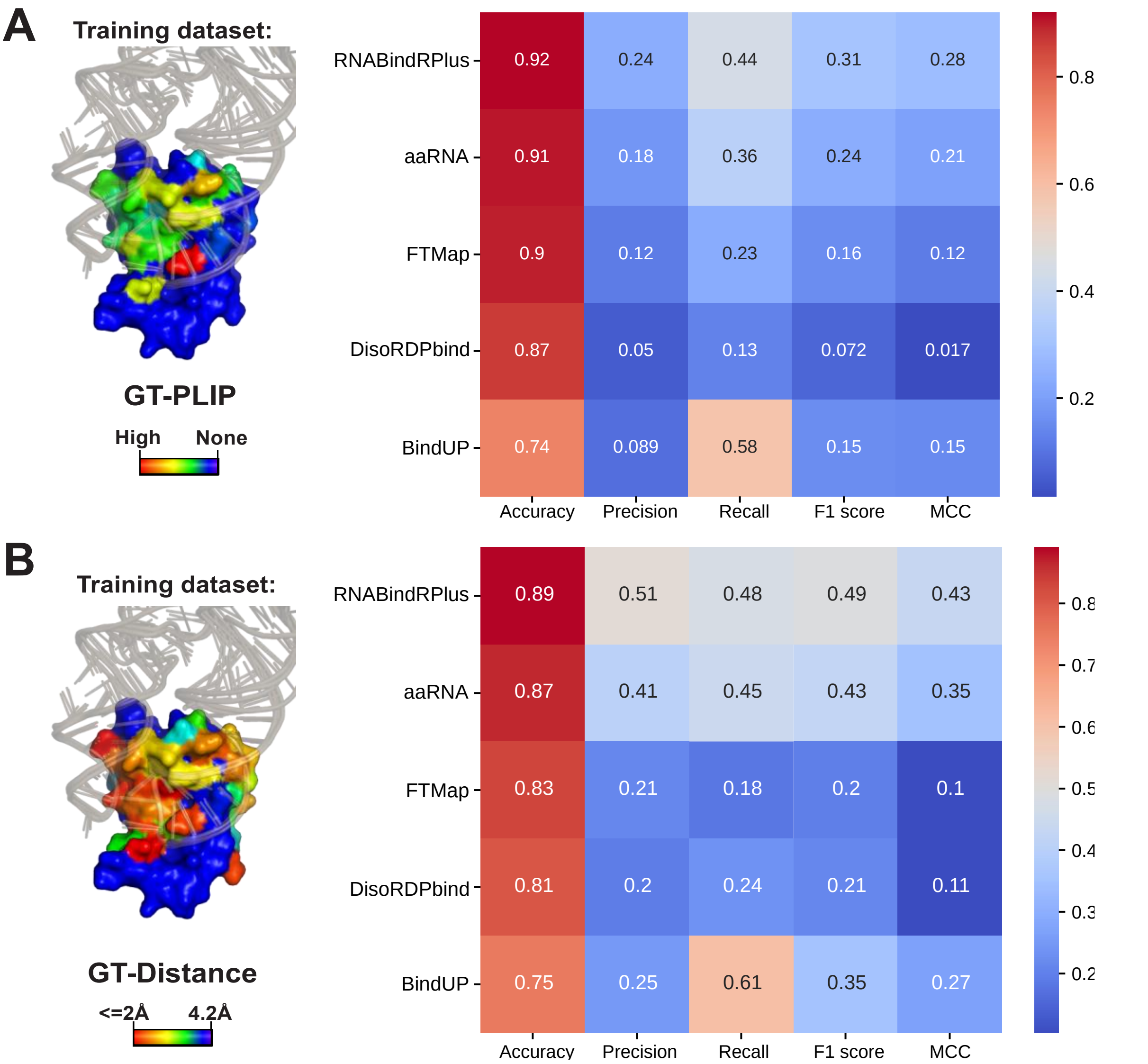


**Figure EV3: Performance metrics for protein-RNA interaction predictors employed by pyRBDome.**

This heatmap represents the comparative analysis of tools used for predicting amino acid - RNA (aaRNA, RNABindRPlus, DisoRDPbind and BindUP) or small molecule interaction sites (FTMap). TP = True Positives, FP = False Positives, TN = True Negatives, FN = False Negatives. Five key performance metrics were calculated, with higher values indicating higher performance: Accuracy: The proportion of true results (both true positives and true negatives) among the total number examined (TP + TN) / (TP + FP + FN + TN). Precision: Proportion of true positives amongst the total predicted as positive (TP / (TP + FP). Recall (Sensitivity): Proportion of positives correctly identified (TP / (TP + FN). F1 Score: Harmonic mean of precision and recall: (2 * (Precision * Recall) / (Precision + Recall) ). Matthews Correlation Coefficient (MCC): A measure of the quality of the classifications that takes into consideration all four confusion matrix categories (TP, FP, FN and TN): ((TP * TN - FP * FN) / sqrt((TP + FP) * (TP + FN) * (TN + FP) * (TN + FN)).

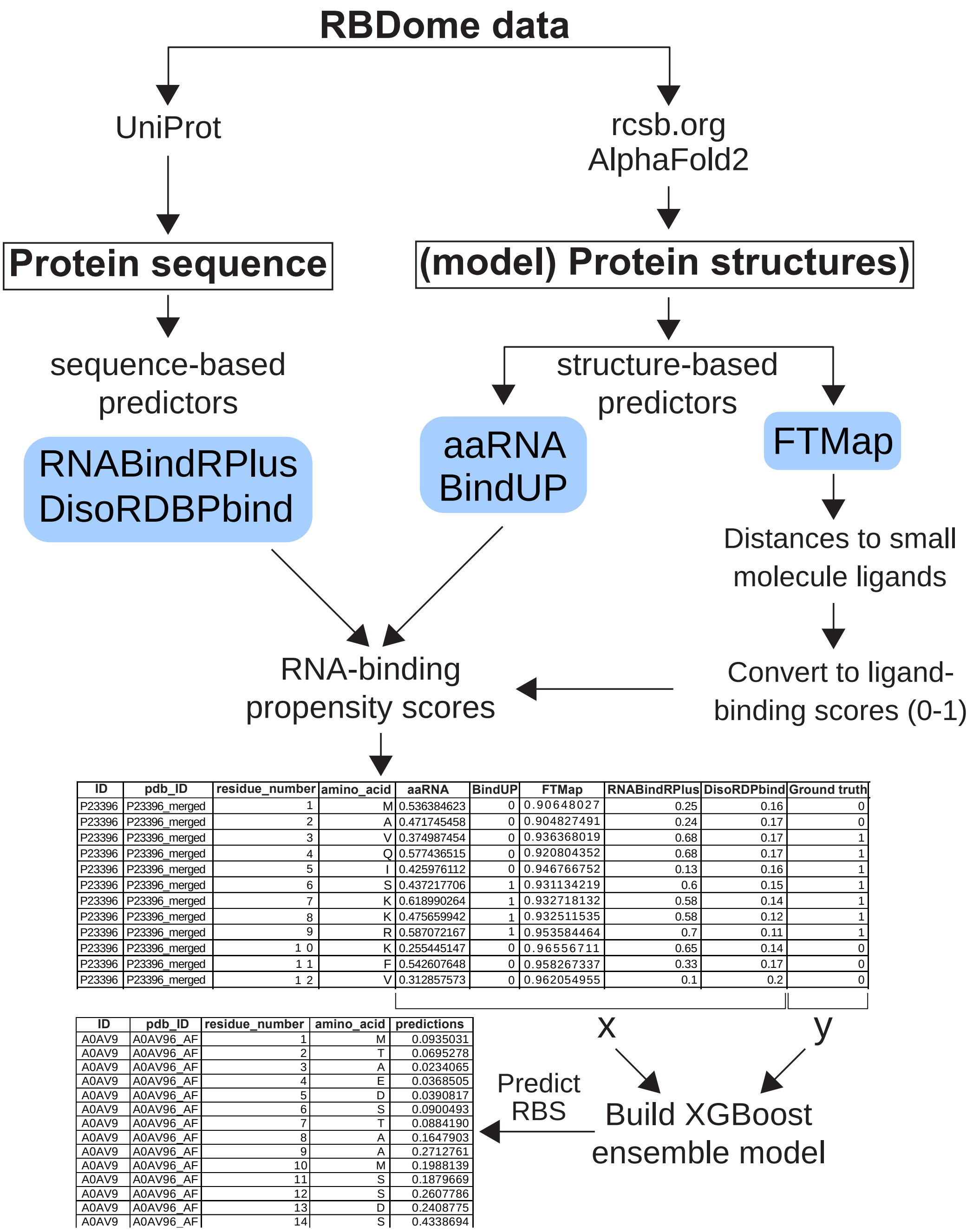


| ID | pdb_ID | residue_number | amino_acid | aaRNA | BindUP | FTMap | RNABindRPlus | DisoRDPbind | Ground truth |
|---|---|---|---|---|---|---|---|---|---|
| P23396 | P23396_merged | 1 | M | 0.536384623 | 0 | 0.90648027 | 0.25 | 0.16 | 0 |
| P23396 | P23396_merged | 2 | A | 0.471745458 | 0 | 0.904827491 | 0.24 | 0.17 | 0 |
| P23396 | P23396_merged | 3 | V | 0.374987454 | 0 | 0.936368019 | 0.68 | 0.17 | 1 |
| P23396 | P23396_merged | 4 | Q | 0.577436515 | 0 | 0.920804352 | 0.68 | 0.17 | 1 |
| P23396 | P23396_merged | 5 | I | 0.425976112 | 0 | 0.946766752 | 0.13 | 0.16 | 1 |
| P23396 | P23396_merged | 6 | S | 0.437217706 | 1 | 0.931134219 | 0.6 | 0.15 | 1 |
| P23396 | P23396_merged | 7 | K | 0.618990264 | 1 | 0.932718132 | 0.58 | 0.14 | 1 |
| P23396 | P23396_merged | 8 | K | 0.475659942 | 1 | 0.932511535 | 0.58 | 0.12 | 1 |
| P23396 | P23396_merged | 9 | R | 0.587072167 | 1 | 0.953584464 | 0.7 | 0.11 | 1 |
| P23396 | P23396_merged | 1 0 | K | 0.255445147 | 0 | 0.96556711 | 0.65 | 0.14 | 0 |
| P23396 | P23396_merged | 1 1 | F | 0.542607648 | 0 | 0.958267337 | 0.33 | 0.17 | 0 |
| P23396 | P23396_merged | 1 2 | V | 0.312857573 | 0 | 0.962054955 | 0.1 | 0.2 | 0 |

| ID | pdb_ID | residue_number | amino_acid | predictions |
|---|---|---|---|---|
| A0AV9 | A0AV96_AF | 1 | M | 0.0935031 |
| A0AV9 | A0AV96_AF | 2 | T | 0.0695278 |
| A0AV9 | A0AV96_AF | 3 | A | 0.0234065 |
| A0AV9 | A0AV96_AF | 4 | E | 0.0368505 |
| A0AV9 | A0AV96_AF | 5 | D | 0.0390817 |
| A0AV9 | A0AV96_AF | 6 | S | 0.0900493 |
| A0AV9 | A0AV96_AF | 7 | T | 0.0884190 |
| A0AV9 | A0AV96_AF | 8 | A | 0.1647903 |
| A0AV9 | A0AV96_AF | 9 | A | 0.2712761 |
| A0AV9 | A0AV96_AF | 10 | M | 0.1988139 |
| A0AV9 | A0AV96_AF | 11 | S | 0.1879669 |
| A0AV9 | A0AV96_AF | 12 | S | 0.2607786 |
| A0AV9 | A0AV96_AF | 13 | D | 0.2408775 |
| A0AV9 | A0AV96_AF | 14 | S | 0.4338694 |

**Figure EV4: Schematic representation of how the data from the aaRNA, BindUP, RNABindRPlus, FTMap and DisoRDPbind prediction results are used to train the XGBoost models.** Data from those tools that provide RNA-binding propensity values between 0 and 1 is directly fed to XGBoost. The x and y characters indicate the experimental and ground truth data, respectively. Values for BindUP, with 10 or higher indicating an RNA-binding site, were normalised to values between 0 and 1. To enable analysis of the FTMap docking results with XGBoost, we calculated the minimum distance of each amino acid in the PDB file to docked ligands. These values were then converted to values between 0 and 1, with the highest value indicating a high ligand-binding score. These values were then passed on to XGBoost. Eighty percent of the GT-PLIP and GT-Distance datasets was used for training purposes and 20% for testing. Once the parameters for the models were optimised, they were used to predict RNA-binding amino acids for the proteins in the RBS-ID dataset (column ‘predictions’). These values represented RNA-binding probabilities. All the analysis results are provided in Datasets EV4 and EV5.

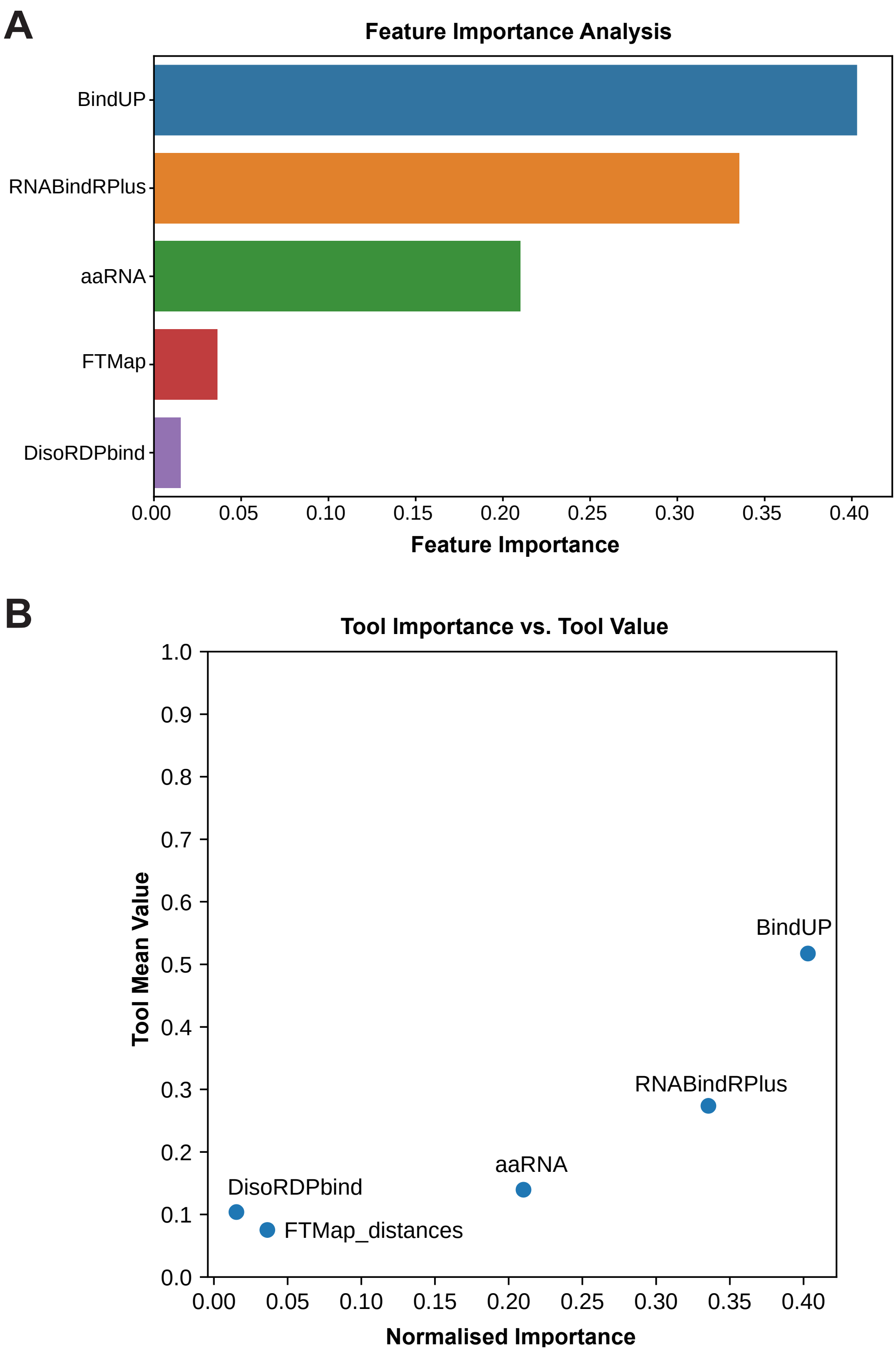


**Figure EV5: Evaluation of predictor significance in XGBoost model efficacy**.
(**A**) The relative importance of different predictors as determined by our XGBoost model. The importance is measured based on how much each predictor contributes to the accuracy of the model. The predictors are listed on the y-axis and their corresponding importance on the x-axis.
(**B**) The normalised feature importance of each predictor against its total mean value from the predictions. The x-axis represents the normalised importance assigned by the XGBoost model, while the y-axis shows the mean value of the prediction results from each tool. The mean is calculated using the accumulation of the impurity decrease within each tree, which is essentially the average of how much the decision made in each tree of the XGBoost model helps to improve to the decision making across all trees.

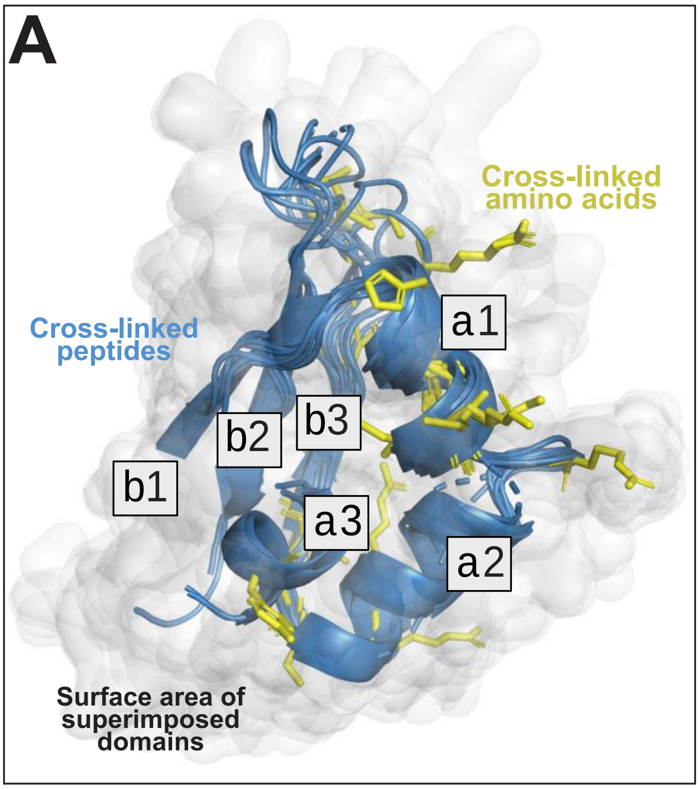


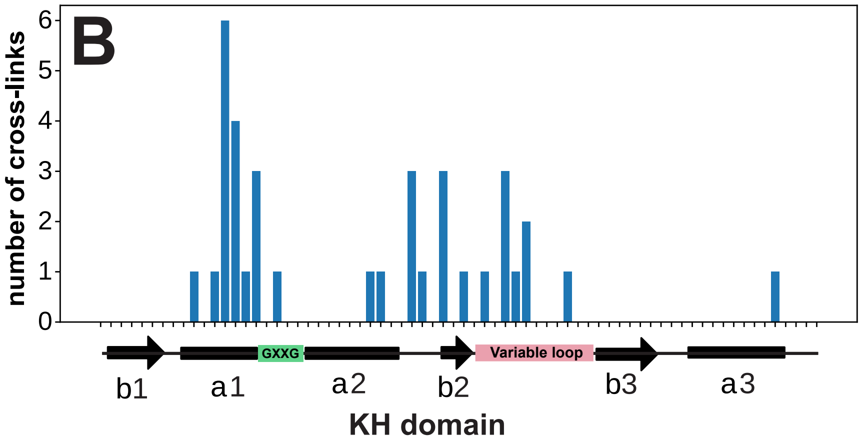


**Figure EV6: Insights into RNA-binding interfaces in protein domains through aggregated amino acid UV cross-linking data can only be generated with data containing many cross-links.**
This figure presents the findings for proteins with KH domains.
(**A**) Superimposed peptide sequences mapped to RRM domains in proteins identified in the RBS-ID dataset. These sequences were aligned on available structural models of RRM domain-containing proteins. The various $\alpha$ and $\beta$ secondary structural elements within the RRM domains are also indicated. Side chains of UV cross-linked amino acids within the domains are highlighted as yellow sticks. The white cloud represents the surface area of the RRM domains.
(**B**) The number of UV cross-links detected in all KH domains at specific positions (y-axis), correlating to their specific positions within the domain (x-axis). Below the x-axis, the consensus secondary structure for KH domains is depicted for reference. GXXG (green) and "variable loop" indicate key regions involved in RNA recognition.

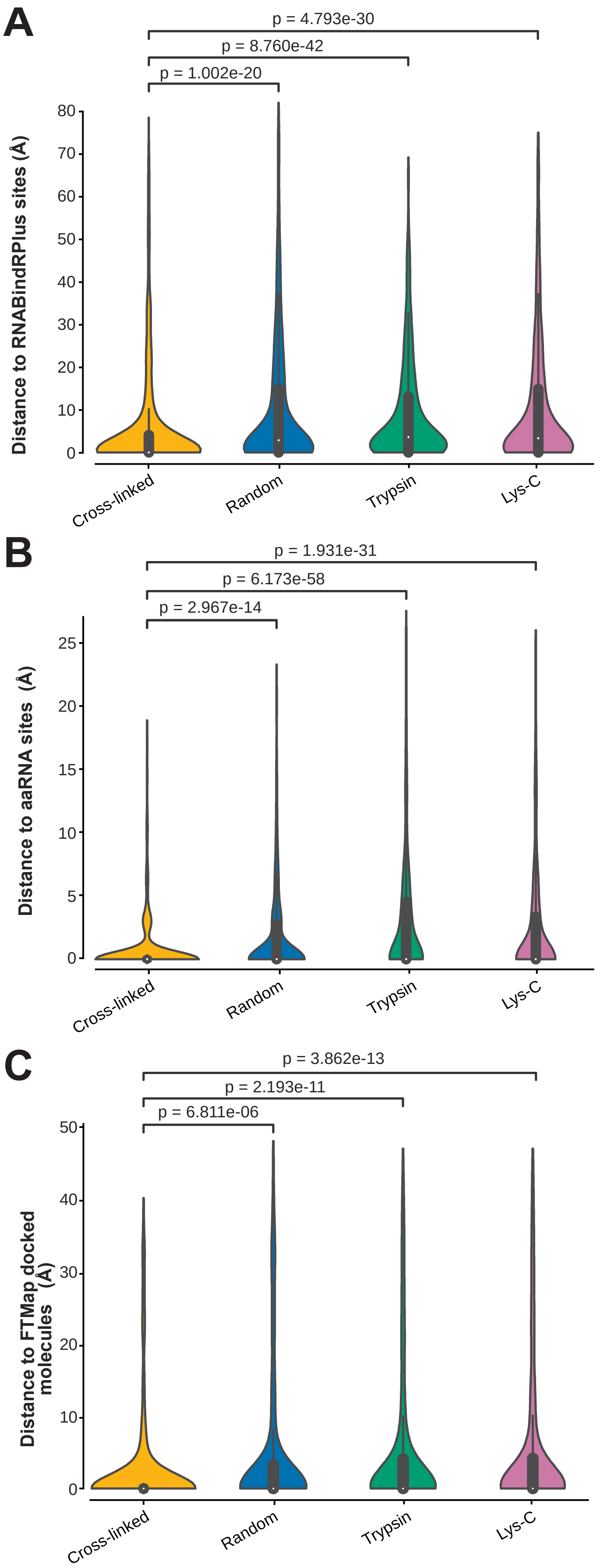


**Figure EV7:** UV cross-linked peptides in RBS-ID data are enriched for RNA and small molecule binding sites or are more likely to be in closer proximity.

(**A**) This panel displays the distribution of distances to RNABindRPlus predicted RBSs identified in cross-linked peptide sequences. Comparative control datasets include randomly generated peptides matching the length distribution and peptide libraries produced *in silico* through Lys-C or Trypsin digestion of the analysed RBPs. P-values were calculated using a two-sided Mann-Whitney U test with Bonferroni correction, highlighting significant differences between the groups as denoted above each comparison. The violins depict density estimates of the distances, where broader sections suggest a higher frequency of amino acids at specific distances. The white dot at the centre of each violin plot signifies the median distance, while the thick bars within the violins indicate the interquartile ranges.

(**B**) As (A) but presenting results for aaRNA predictions.

(**C**) As (A) but illustrating the results for distances to FTMap-docked small molecules.

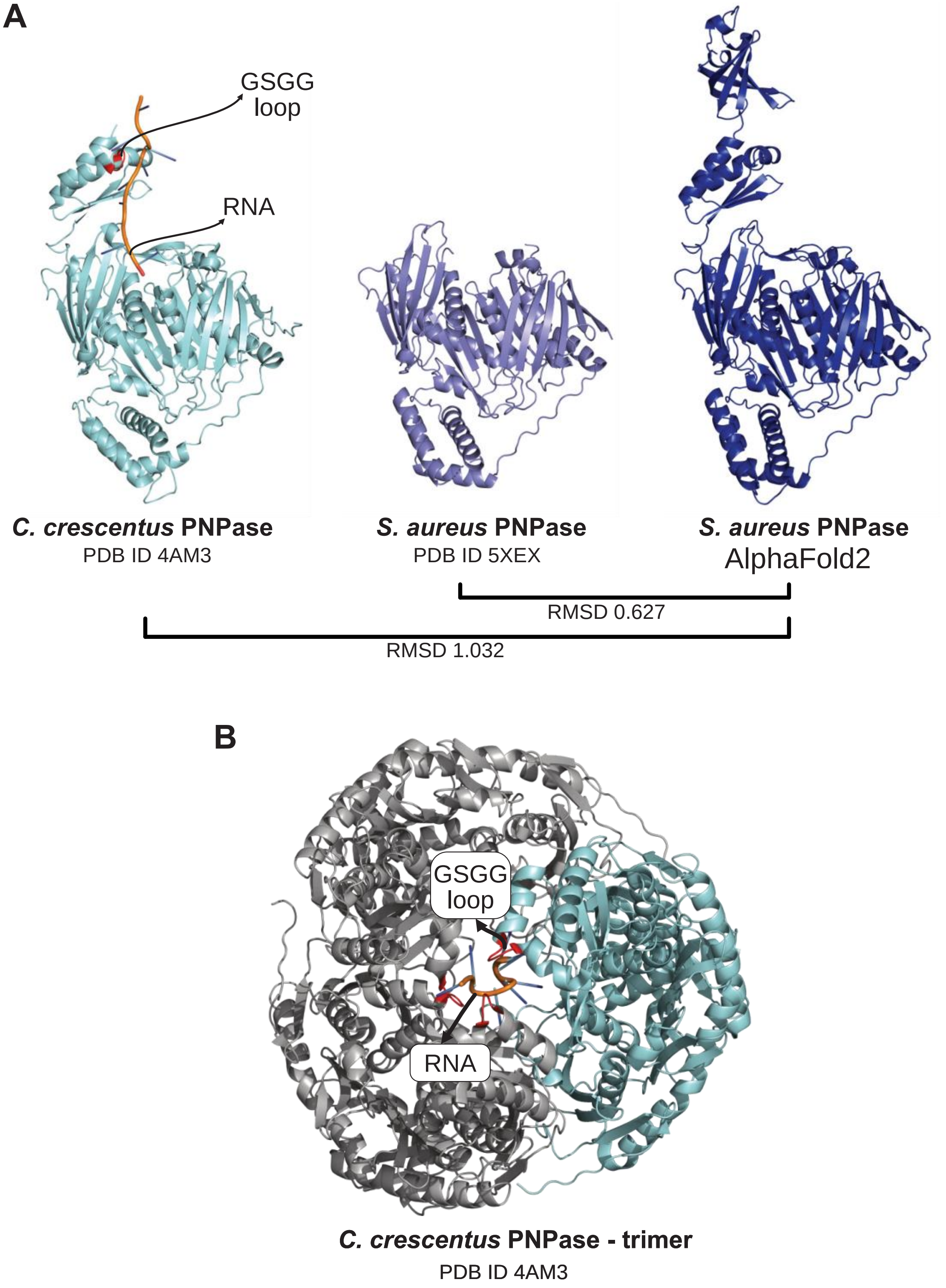


**Figure EV8: PNPase AlphaFold2 model similarity to published crystal structures.**
(**A**) Crystal structure of the *C. crescentus* PNPase trimer bound with RNA (PDB ID 4AM3; (Hardwick *et al*, 2012)). Noted are the positions of the co-crystallised RNA fragment and the RNA-binding GSGG loop.
(**B**) The crystal structure of the *C. crescentus* PNPase monomer in complex with RNA. Highlighted are the co-crystallised RNA fragment and the RNA-binding GSGG loop. Additionally, the crystal structure of the *S. aureus* PNPase active site alongside the AlphaFold2 model is presented. Below the models, the root-mean-square deviation (RMSD) values for the various model comparisons are provided.

### Expanded View Dataset Legends

**Dataset EV1:** Dataset EV1: Typical input file required for performing pyRBDome analyses. Shown are the RBS-ID results (Bae *et al*, 2020) that were subjected to pyRBDome analyses. The first column (UniProt) shows the UniProt ID associated with the proteins of interest. The second (Protein) column shows an alternative protein name for the UniProt ID. The RBS_aa column contains the amino acid residues that were reported to cross-link to RNA. The RBS_aa_location column indicates where in the cross-linked peptide (next column) the cross-linked amino acid was located. The Peptide column provides the sequences of the cross-linked peptides.

**Dataset EV2:** An example of a pyRBDome analysis output file. This table displays all the results of the individual tools for each peptide that was analyzed. The first column ('ID') contains the UniProt ID associated with the cross-linked peptides. The second column ('pdb_id') contains the name of the PDB file used for the analyses. The naming convention for these files is as follows: O00425_6GX6, where O00425 is the UniProt ID for the structure, and 6GX6 is the RCSB.org ID. If the name ends with '_AF', an AlphaFold2 model was downloaded for that protein. The 'Found_peptide' column indicates where in the PDB file (as noted in the 'pdb_id' column) the peptide sequence was mapped. Consider the following peptide: 'GRLLGVCCSVDNC'. This was mapped to pdb_id A0AV96_AF, resulting in: 139_A_grllgvccsvdnc_151_A. This means that the peptide starts at residue number 139 and ends at 151 in chain A of the PDB file. The peptide sequence is now in lowercase. The columns 'BindUP_results', 'aaRNA_results', 'DisoRDPbind_results', 'FTMap_results', and 'RNABindRPlus_results' show which amino acids in the peptide sequences are predicted to bind RNA by these tools. For example, consider the aaRNA results from the previous peptide example: 139_A_GRllgvccsvdnc_151_A. Here, the 'G' and 'R' are in uppercase, indicating that aaRNA predicts these two amino acids in the peptide sequence bind RNA. An example of an FTMap prediction result is: 139_A_grllgvCCSVDNC_151_A, where the uppercase amino acids are those within 4.2Å of small molecules docked onto the PDB file by FTMap. The columns ending with '_distances' show the shortest distance (in Å) of a peptide to the nearest predicted RNA-binding site. A 0 indicates that the predicted RNA-binding amino acid is located in the peptide sequence. In some rows, you will see 'not_found' or 'no_data'. 'Not_found' indicates that the peptide could not be mapped to the structure, possibly because the PDB file did not contain the complete protein sequences, and therefore the peptide/amino acid could not be mapped. This should not be a problem with AlphaFold2 models. 'No data' indicates that either the peptide was not mapped to the PDB file structure, or the analysis with one of the predictors failed and thus did not return a result.

**Dataset EV3:** The same as for dataset EV2, but now for the cross-linked amino acids that were reported in the RBS-ID data.

**Dataset EV4:** Table showing the pyRBDome prediction results and all the experimental data available for the proteins used to generate our ground truth datasets. Each row in this table represents the results for a single amino acid from a specific protein. The first column ('ID') contains the UniProt ID associated with the cross-linked peptides. The second column ('pdb_id') contains the name of the PDB file used for the analyses. The 'residue_number' column shows the residue number for that amino acid from the PDB file that was analyzed. The 'aaRNA', 'BindUP', 'RNABindRPlus', and 'DisoRDPbind' columns display the RNA-binding probabilities returned by these prediction algorithms for each amino acid (values between 0-1). The 'FTMap_distances' column shows the distances (in Å) of the amino acid to the nearest small molecule docked on the structure by FTMap. The 'Domains' column highlights which amino acids are part of a domain (indicated by a '1'). The 'Peptide' column shows which amino acids were detected in cross-linked peptides (indicated by a '1'). The 'Cross-linked_amino_acid' column highlights the cross-linked amino acid that was detected in the protein (indicated by a '1'). The ‘predictions’ column shows the RNA-binding probabilities determined by our XGBoost model trained on the GT-Distance ground truth data.

**Dataset EV5:** Table showing results from the analysis of the protein structures used to generate the GT-PLIP and GT-Distance ground truth datasets. The first column ('ID') contains the UniProt ID associated with the cross-linked peptides. The second column ('pdb_id') contains the name of the PDB file used for the analyses. The 'residue_number' column shows the residue number for that amino acid from the PDB file that was analyzed. The term 'merged' in the file name indicates that this PDB file was created by consolidating RNA distance analysis results from multiple PDB files, with the shortest distance to RNA indicated in the b-factor column. The 'Peptide' column indicates which amino acids were detected in cross-linked peptides (marked by a '1'). The 'Cross-linked_amino_acid' column highlights the specific cross-linked amino acid detected in the protein (also marked by a '1'). The ‘Distance_to_RNA’ column shows the minimal distance (in Å) of the amino acid to the nearest RNA atom. The ‘PLIP analysis results’ columns show the frequency of hydrogen bonds, hydrophobic interactions, salt bridges, $\pi$-cation, and $\pi$-stacking interactions detected by PLIP (Protein-Ligand Interaction Profiler; (Adasme *et al*, 2021)) in the associated PDB file. The ‘all bonds’ column is a sum of all these interaction types detected. The ‘Distance_to_PLIP’ column indicates the distance (in Å) of an amino acid to the nearest RNA-binding amino acid as determined by PLIP.
